## Supplementary Figures for "Ribosome-binding protein 1, RRBP1, maintains peroxisome biogenesis"

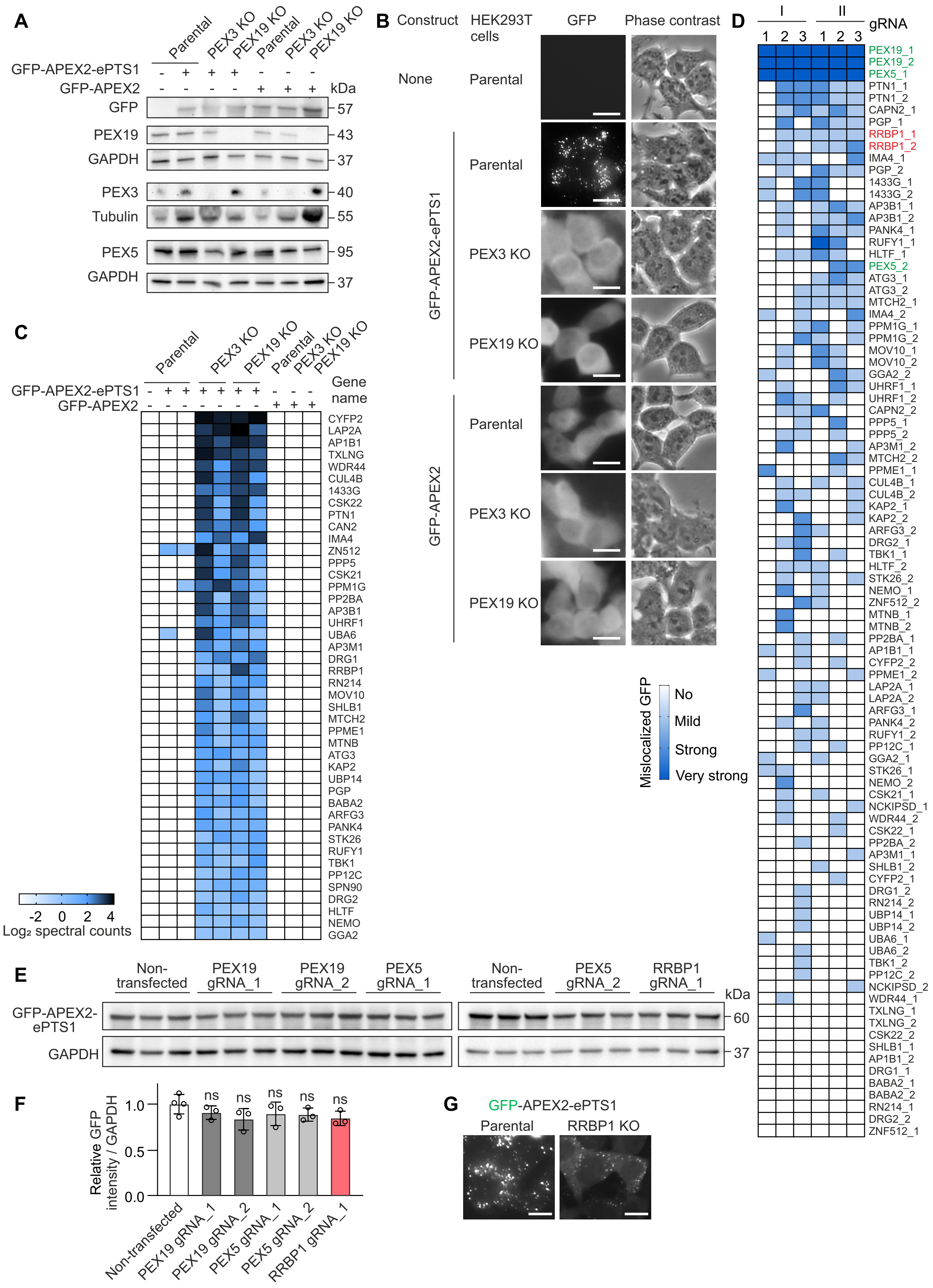

A
Parental
PEX3 KO
PEX19 KO
GFP-APEX2-ePTS1
GFP-APEX2
kDa
GFP
PEX19
GAPDH
PEX3
Tubulin
PEX5
57
43
37
40
55
95
B
Construct
HEK293T cells
GFP
Phase contrast
None
GFP-APEX2-ePTS1
GFP-APEX2
C
Gene name
CYFP2
LAP2A
AP1B1
TXLNG
WDR44
CUL4B
1433G
CSK22
PTN1
CAN2
IMA4
ZN512
PPP5
CSK21
PPM1G
PP2BA
AP3B1
UHRF1
UBA6
AP3M1
DRG1
RRBP1
RN214
MOV10
SHLB1
MTCH2
PPME1
MTNB
ATG3
KAP2
UBP14
PGP
BABA2
ARFG3
PANK4
STK26
RUFY1
TBK1
PP12C
SPN90
DRG2
HLTF
NEMO
GGA2
-2 0 2 4
Log₂ spectral counts
D
I
II
gRNA
1 2 3 1 2 3
PEX19_1
PEX19_2
PEX5_1
PTN1_1
PTN1_2
CAPN2_1
PGP_1
RRBP1_1
RRBP1_2
IMA4_1
PGP_2
1433G_1
1433G_2
AP3B1_1
AP3B1_2
PANK4_1
RUFY1_1
HLTF_1
PEX5_2
ATG3_1
ATG3_2
MTCH2_1
IMA4_2
PPM1G_1
PPM1G_2
MOV10_1
MOV10_2
GGA2_2
UHRF1_1
UHRF1_2
CAPN2_2
PPP5_1
PPP5_2
AP3M1_2
MTCH2_2
PPME1_1
CUL4B_1
CUL4B_2
KAP2_1
KAP2_2
ARFG3_2
DRG2_1
TBK1_1
HLTF_2
STK26_2
NEMO_1
ZNF512_2
MTNB_1
MTNB_2
PP2BA_1
AP1B1_1
CYFP2_2
PPME1_2
LAP2A_1
LAP2A_2
ARFG3_1
PANK4_2
RUFY1_2
PP12C_1
GGA2_1
STK26_1
NEMO_2
CSK21_1
NCKIPSD_1
WDR44_2
CSK22_1
PP2BA_2
AP3M1_1
SHLB1_2
CYFP2_1
DRG1_2
RN214_2
UBP14_1
UBP14_2
UBA6_1
UBA6_2
TBK1_2
PP12C_2
NCKIPSD_2
WDR44_1
TXLNG_1
TXLNG_2
CSK22_2
SHLB1_1
AP1B1_2
DRG1_1
BABA2_1
BABA2_2
RN214_1
DRG2_2
ZNF512_1
Mislocalized GFP
No
Mild
Strong
Very strong
E
Non-transfected
PEX19 gRNA_1
PEX19 gRNA_2
PEX5 gRNA_1
PEX5 gRNA_2
RRBP1 gRNA_1
GFP-APEX2-ePTS1
GAPDH
60
37
F
Relative GFP intensity / GAPDH
1.0
0.5
0.0
ns
Non-transfected
PEX19 gRNA_1
PEX19 gRNA_2
PEX5 gRNA_1
PEX5 gRNA_2
RRBP1 gRNA_1
G
GFP-APEX2-ePTS1
Parental
RRBP1 KO

**Supplementary Figure 1. Systematic identification of peroxisome biogenesis factors using proximity labeling and CRISPR/Cas9 microscopy-based screen.**

(A) Western blotting analysis of HEK293T clonal cell lines stably expressing GFP-APEX2-ePTS1 or GFP-APEX2 constructs used for APEX2-based proximity labeling (Fig. 1A). GAPDH or tubulin was used as a loading control. (B) Live-cell fluorescence microscopy images showing intracellular localization of GFP-APEX2-ePTS1 or GFP-APEX2 constructs in HEK293T cells used for APEX2-based proximity labeling. The localization of GFP-containing constructs was visualized using green channel. Scale bar, 20 μm. (C) Heat map of the potential peroxisome biogenesis factors identified by APEX2-based proximity labeling (Fig. 1B). (D) Heat map of CRISPR/Cas9 microscopy assessing the phenotype of HEK293T cells. HEK293T cells stably expressing GFP-APEX2-ePTS1 were transiently transfected with gRNAs and Cas9-mCherry plasmid as shown in (A). Two rounds of transfection were performed (I and II). Each round the phenotype of the cells was assessed three times at 7, 10 and 14 post-transfection days using live-cell fluorescence microscopy. Cells were manually classified based on GFP localization: “no” (GFP localized exclusively to peroxisomes, no mislocalization), “mild” (some mislocalized GFP), “strong” (substantial mislocalized GFP), and “very strong” (extensive mislocalized GFP). gRNAs targeting PEX5 or PEX19 were used as positive controls (in green). (E) Western blot analysis of GFP-APEX2-ePTS protein level in HEK293T cells stably expressing GFP-APEX2-ePTS1 before and after 13 days of transfection with Cas9-mCherry plasmid and gRNA targeting PEX19, PEX5 or RRBP1. Anti-GFP antibody was used to detect the GFP-APEX2-ePTS protein. GAPDH was used as a loading control. (F) Quantification of western blot images represented in (E) (n = 3). Values are normalized to the non-transfected cells, set to 1. In all graphs data are presented as mean ± SD; ns, not significant as compared to the non-transfected cells (unpaired t-tests). (G) Live-cell fluorescence microscopy images showing intracellular localization of stability ex-pressing GFP-APEX2-ePTS1 construct in RRBP1 KO or parental HEK293T cells. The localization of GFP-containing constructs was visualized using green channel. Scale bar, 10 μm.

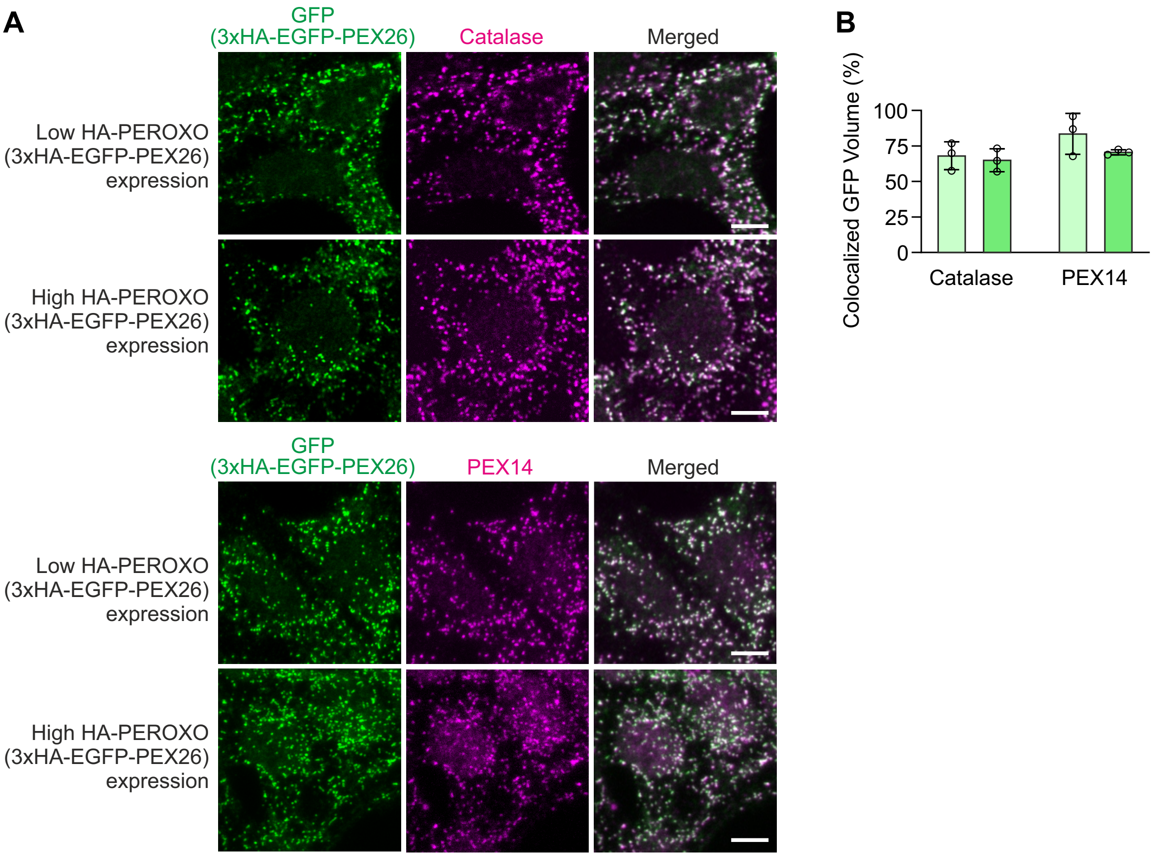


**Supplementary Figure 2. HA-PEROXO (3×HA-EGFP-PEX26) construct colocalizes with peroxisomal markers.**

(A) Confocal microscopy images of HEK293T cells stably expressing 3×HA-EGFP-PEX26 (HA-PEROXO). Immunostaining was performed using anti-GFP, anti-catalase or anti-PEX14 antibody. Scale bar, 10 µm. (B) Colocalization analysis of confocal microscopy images represented in (A). The data are presented as mean ± SD.

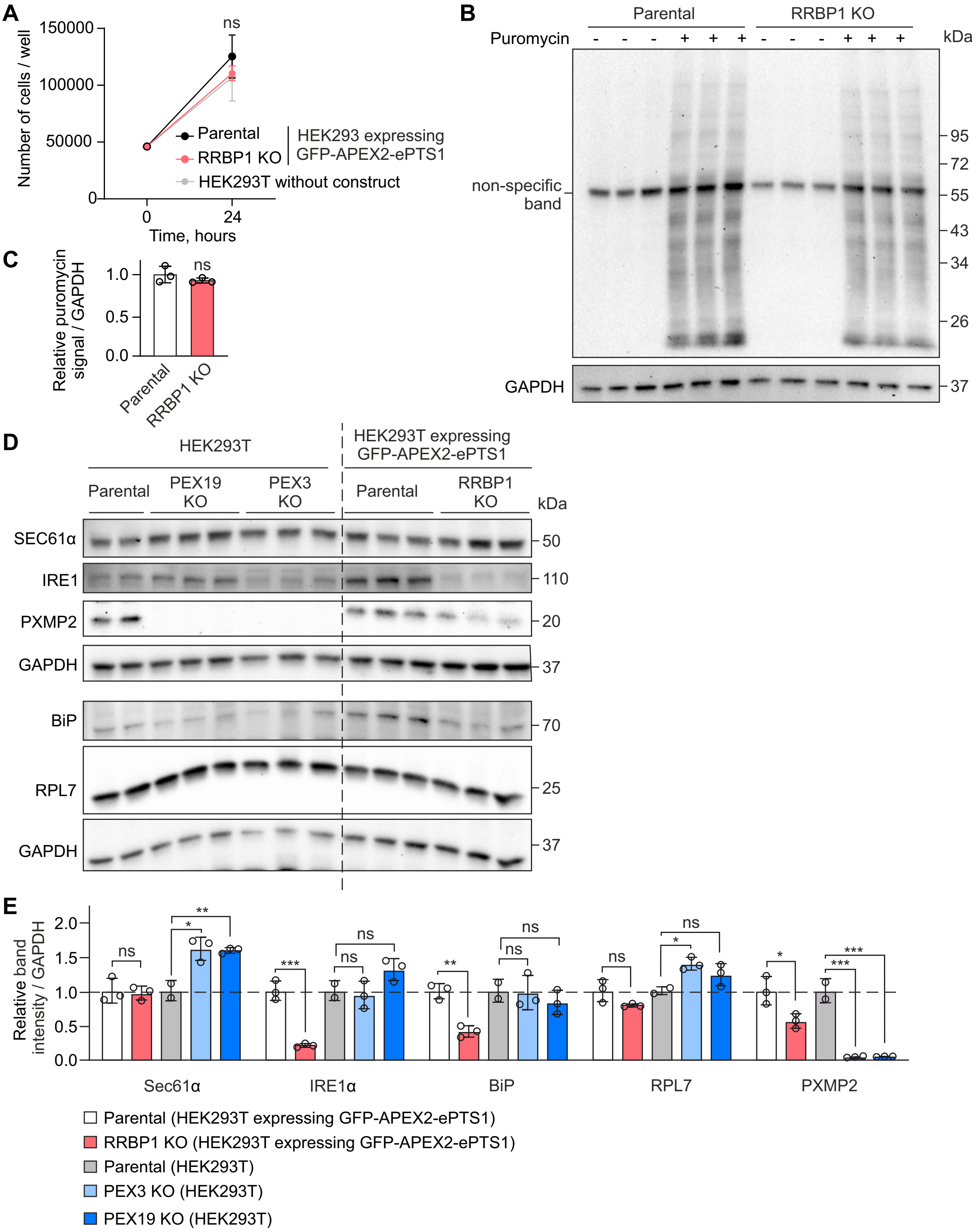

A
Number of cells / well
150000
100000
50000
0
ns
Parental
RRBP1 KO
HEK293 expressing GFP-APEX2-ePTS1
HEK293T without construct
0
24
Time, hours
B
Parental
RRBP1 KO
Puromycin
kDa
95
72
55
43
34
26
non-specific band
GAPDH
37
C
Relative puromycin signal / GAPDH
1.0
0.5
0.0
ns
Parental
RRBP1 KO
D
HEK293T
HEK293T expressing GFP-APEX2-ePTS1
Parental
PEX19 KO
PEX3 KO
Parental
RRBP1 KO
kDa
SEC61α
IRE1
PXMP2
GAPDH
BiP
RPL7
GAPDH
50
110
20
37
70
25
37
E
Relative band intensity / GAPDH
2.0
1.5
1.0
0.5
0.0
Sec61α
IRE1α
BiP
RPL7
PXMP2
Parental (HEK293T expressing GFP-APEX2-ePTS1)
RRBP1 KO (HEK293T expressing GFP-APEX2-ePTS1)
Parental (HEK293T)
PEX3 KO (HEK293T)
PEX19 KO (HEK293T)

**Supplementary Figure 3. RRBP1 KO does not affect cell proliferation and does not induce ER stress.** (A) Proliferation rate of HEK293T cells stably expressing GFP-APEX2-ePTS1 (parental, RRBP1 KO) or HEK293T cells not expressing peroxisomal targeted construct was analyzed using automated cell counter. (B) Protein translation rate in RRBP1 KO or parental HEK293T cells stably expressing GFP-APEX2-ePTS1 was analyzed by puromycin incorporation into nascent proteins. Cells were labeled with puromycin (10 µg/ml, 20 min) and the protein extracts were analyzed by western blotting using anti-puromycin antibody. GAPDH was used as a loading control. (C) Quantification of western blot images represented in (B) (n = 3). (D) Western blot analysis of RRBP1 KO HEK293T cells stably expressing GFP-APEX2-ePTS1. Parental, PEX3 and PEX19 KO HEK293T not expressing peroxisomal targeted construct were used as controls. GAPDH was used as a loading control. (E) Quantification of western blot images represented in (D) (n = 3). Each western blot analysis was performed at least two times. Values are normalized to the corresponding parental cell line, set to 1. In all graphs data are presented as mean ± SD. *P < 0.05, **P < 0.01, ***P 0.001, ns, not significant as compared to the parental cell line (unpaired t-tests).

### Unprocessed western blot images related to Figure 2A

Western blot analysis of RRBP1 KO HEK293T cells stably expressing GFP-APEX2-ePTS1.
Parental, PEX3 and PEX19 KO HEK293 cells without peroxisomal targeted construct were used as controls.
Areas presented in the figure are indicated.

Blot 1: 7.5% SDS PAGE. RRBP1 and GAPDH were detected on the same blot in the folllowing order: RRBP1 first and GFP last.

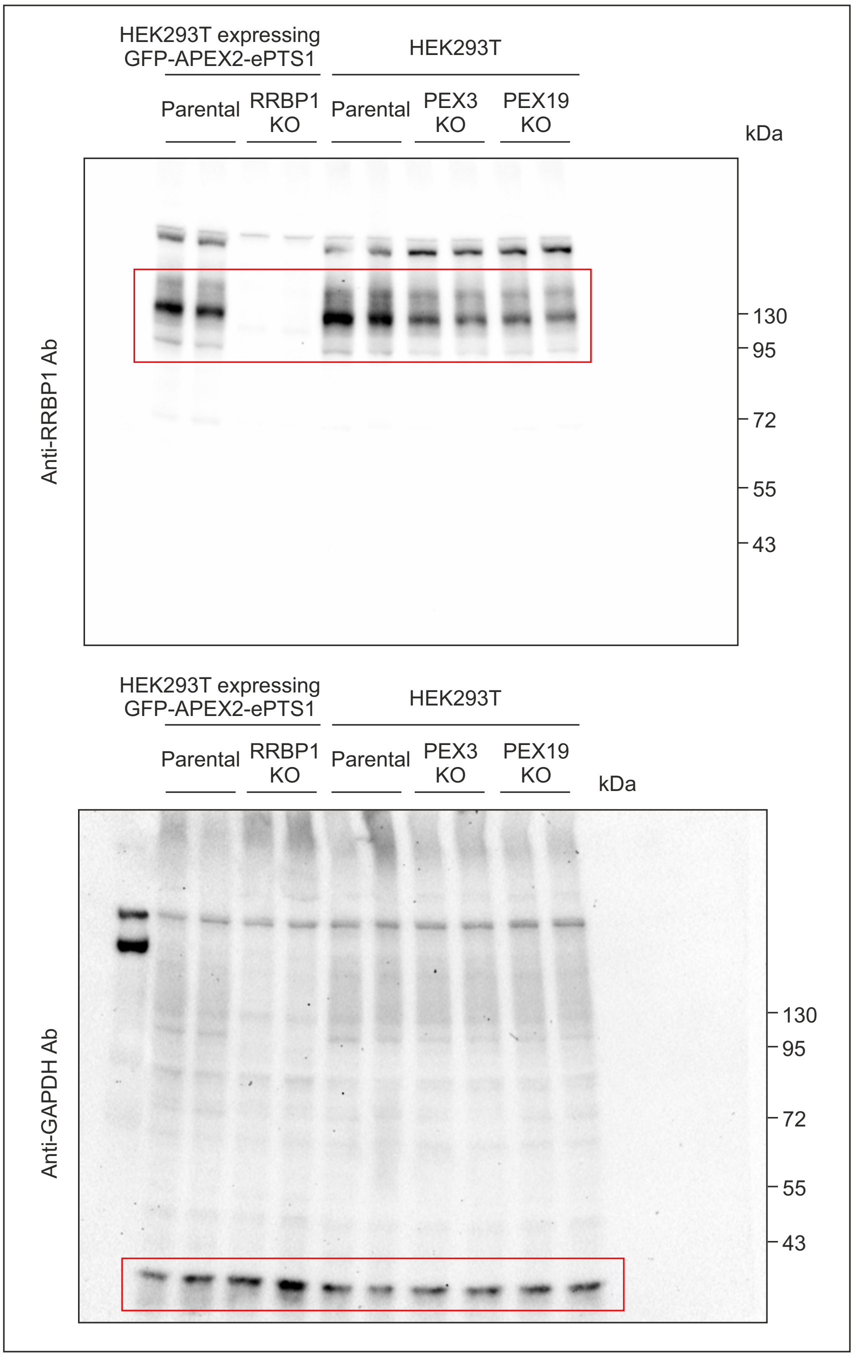

Blot 2: 10% SDS PAGE. PEX5, catalase, TOM20 and GAPDH were detected on the same blot in the folllowing order: catalase, PEX5, TOM20 and GAPDH last.

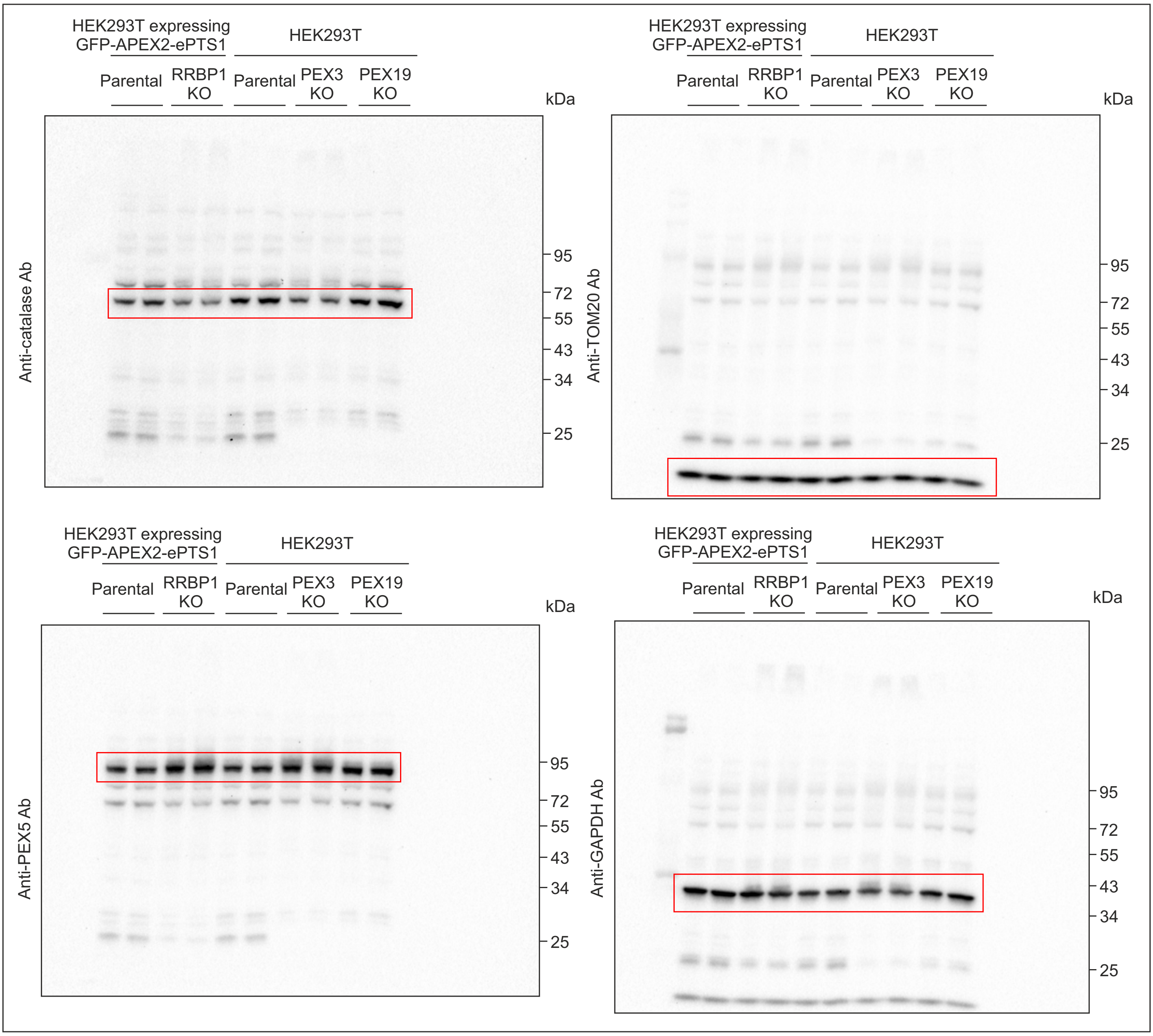

Blot 3: 10% SDS PAGE. ACOX1, PEX14 and GAPDH were detected on the same blot in the folllowing order: ACOX1 first, then GAPDH and PEX14 last.

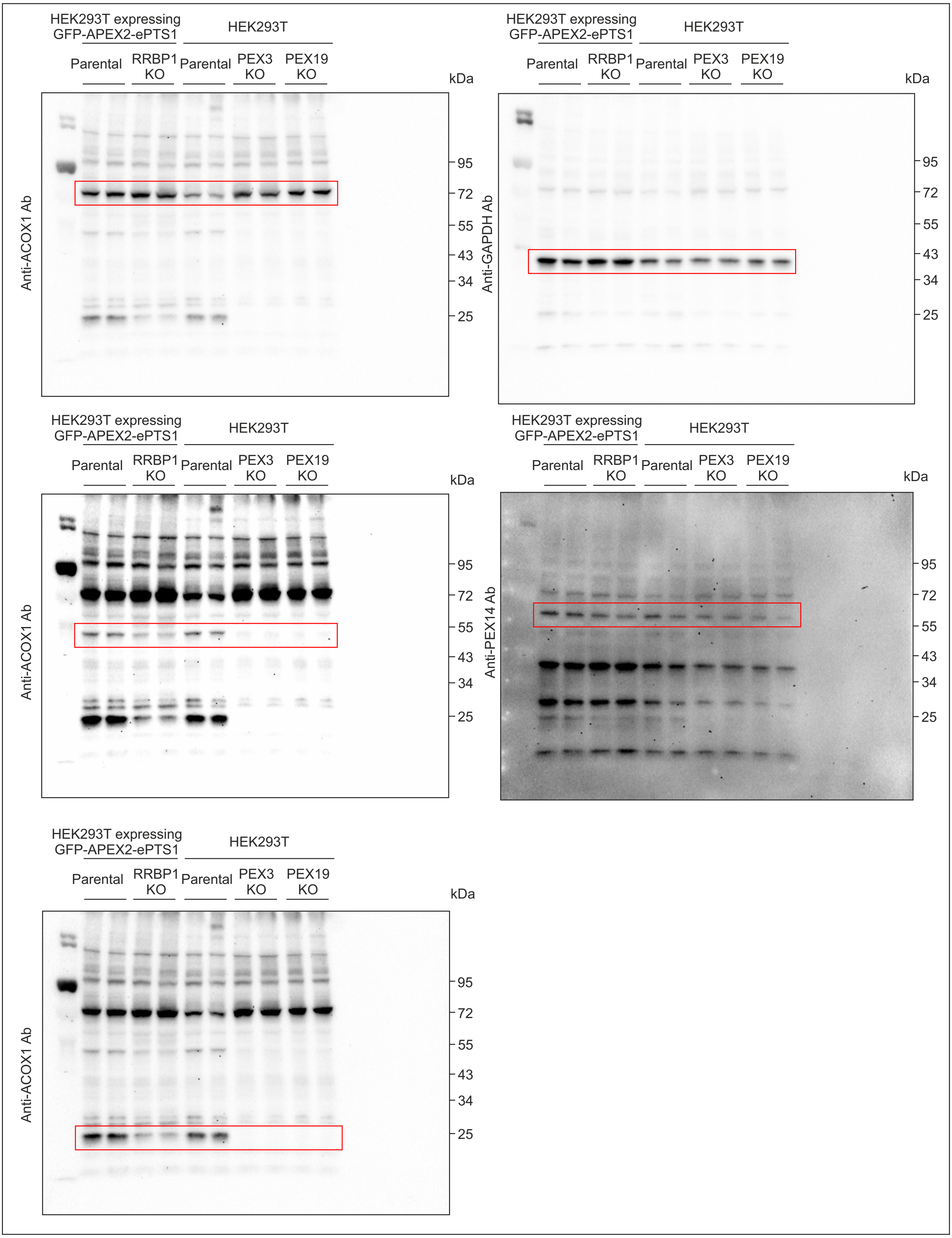

Blot 4: 10% SDS PAGE. PEX3, calnexin, VAPB and GAPDH were detected on the same blot in the folllowing order: calnexin, PEX3, GAPDH and VAPB last.

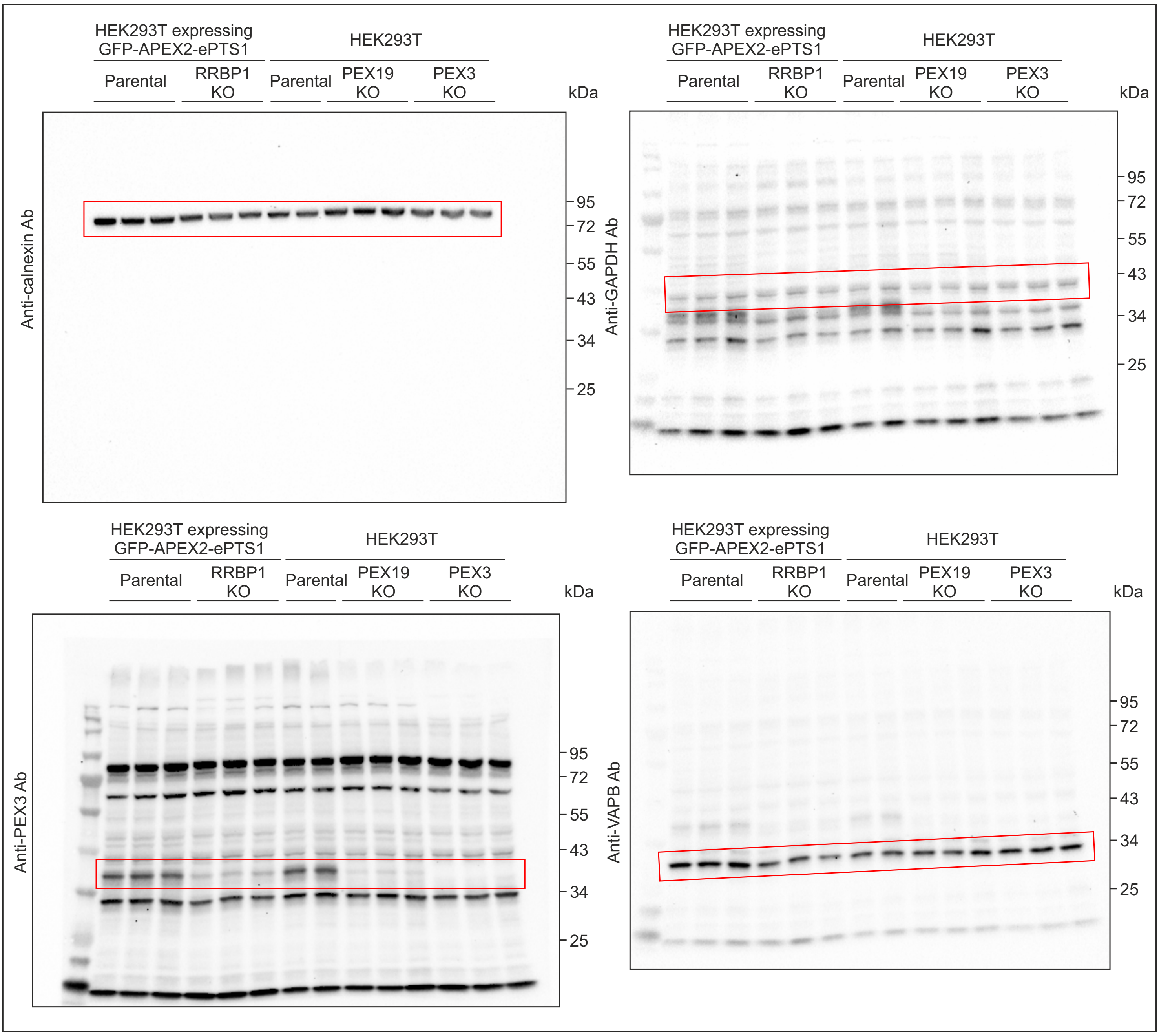

Blot 5: 10% SDS PAGE. PEX19 and GAPDH were detected on the same blot in the folllowing order: PEX19 first and then GAPDH.

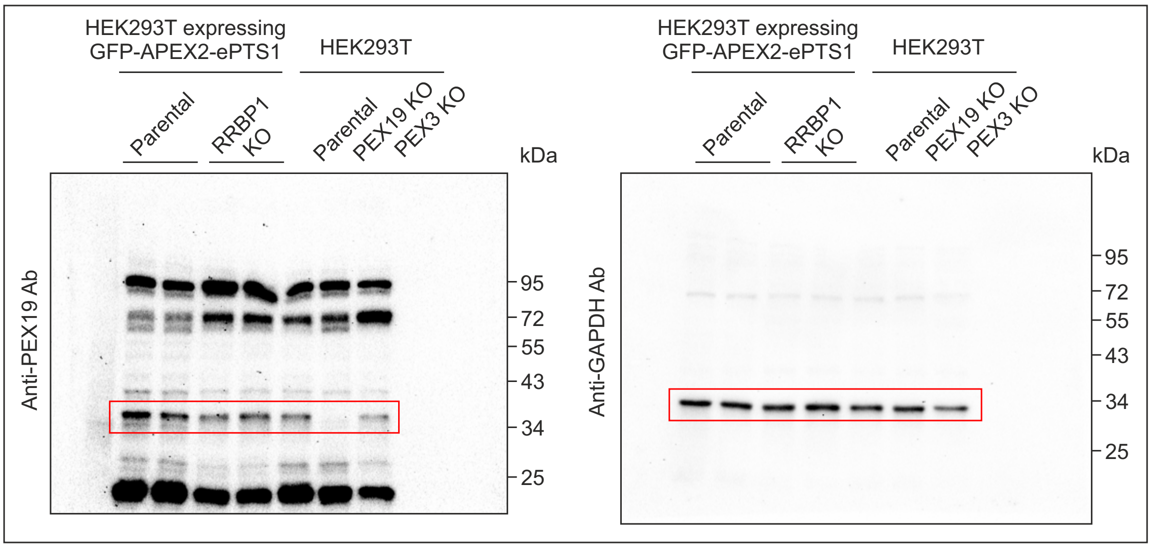


Blot 6: 7.5% SDS PAGE. ACAA1 and GAPDH were detected on the same blot in the folllowing order: ACAA1 first and then GAPDH.

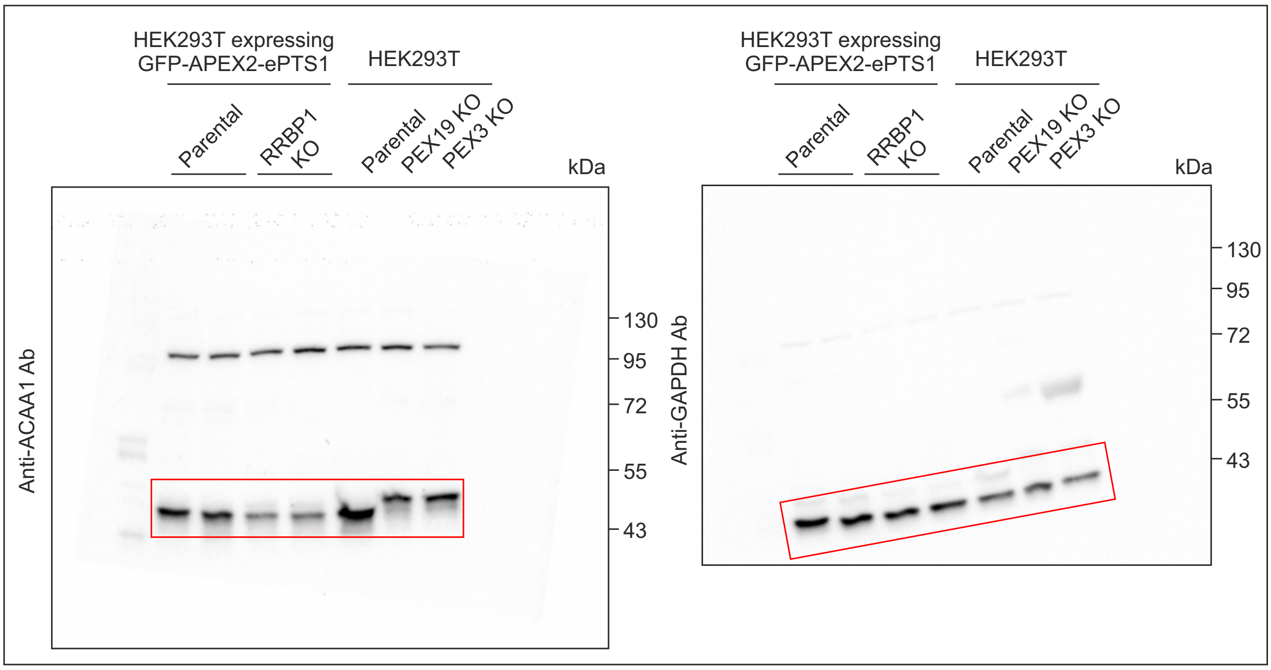

#### Unprocessed western blot images related to Figure 4A

Western blot analysis of RRBP1 silencing in HEK293T cells stably expressing GFP-APEX2-ePTS1.
The cells were transfected with corresponding esiRNAs for 72 h. GFP esiRNA was used as a negative control, esiRNA PEX5 was used as a positive control.
Areas presented in the figure are indicated.

Blot 1: 7.5% SDS PAGE. GFP, RRBP1, PEX5, ACAA1 and GAPDH were detected on the same blot in the folllowing order: RRBP1, ACAA1, PEX4, GFP and GAPDH last.

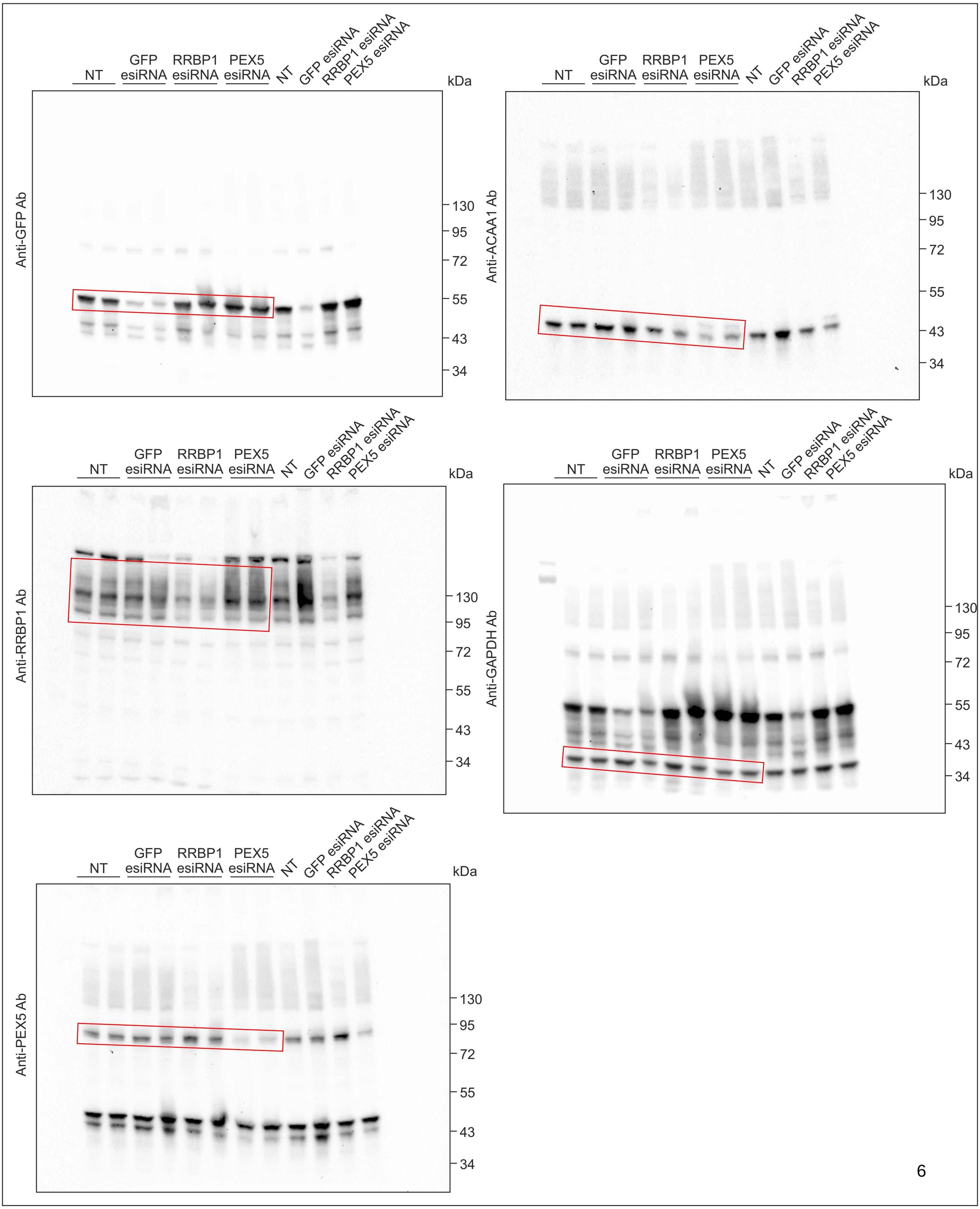

Blot 2: 10% SDS PAGE. ACOX1, catalase and tubulin were detected on the same blot in the folllowing order: ACOX1 first, then catalase and tubulin last.

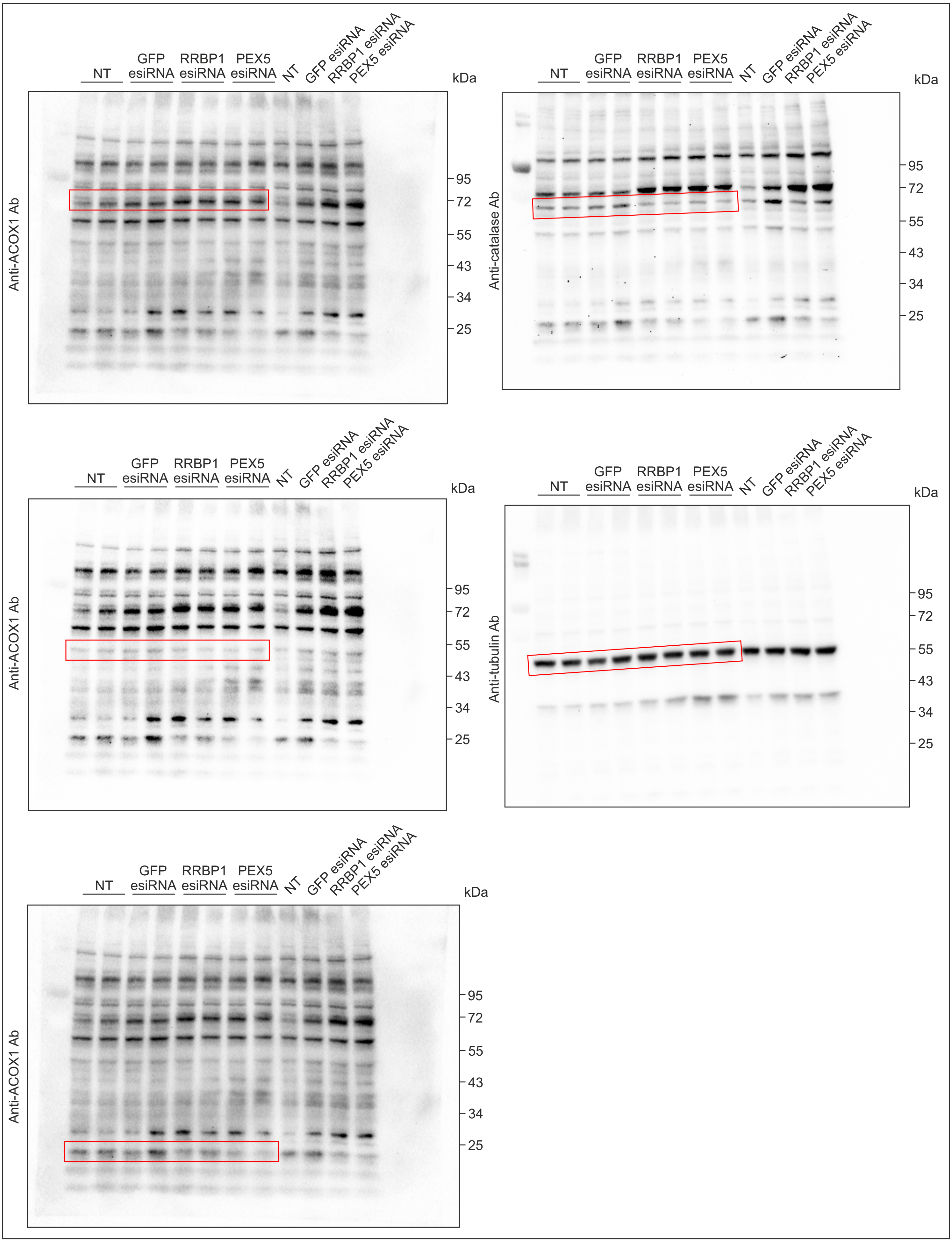

#### Unprocessed western blot images related to Figure 5C

Western blot of HEK293T cells expressing 3×Myc-EGFP-PEX26 (HA-PEROXO) and immunopurified peroxisomes.
Areas presented in the figure are indicated.

Blot 1: 10% SDS PAGE. The blot was cut into four parts. PMP70 and RRBP1 were detected on the upper part of the blot in the folllowing order: RRBP1 first and then PMP70. Catalase was detected on the second part of the blot. GAPDH was detected on the third part of the blot. TOM20 was detected on the lowest part of the blot.

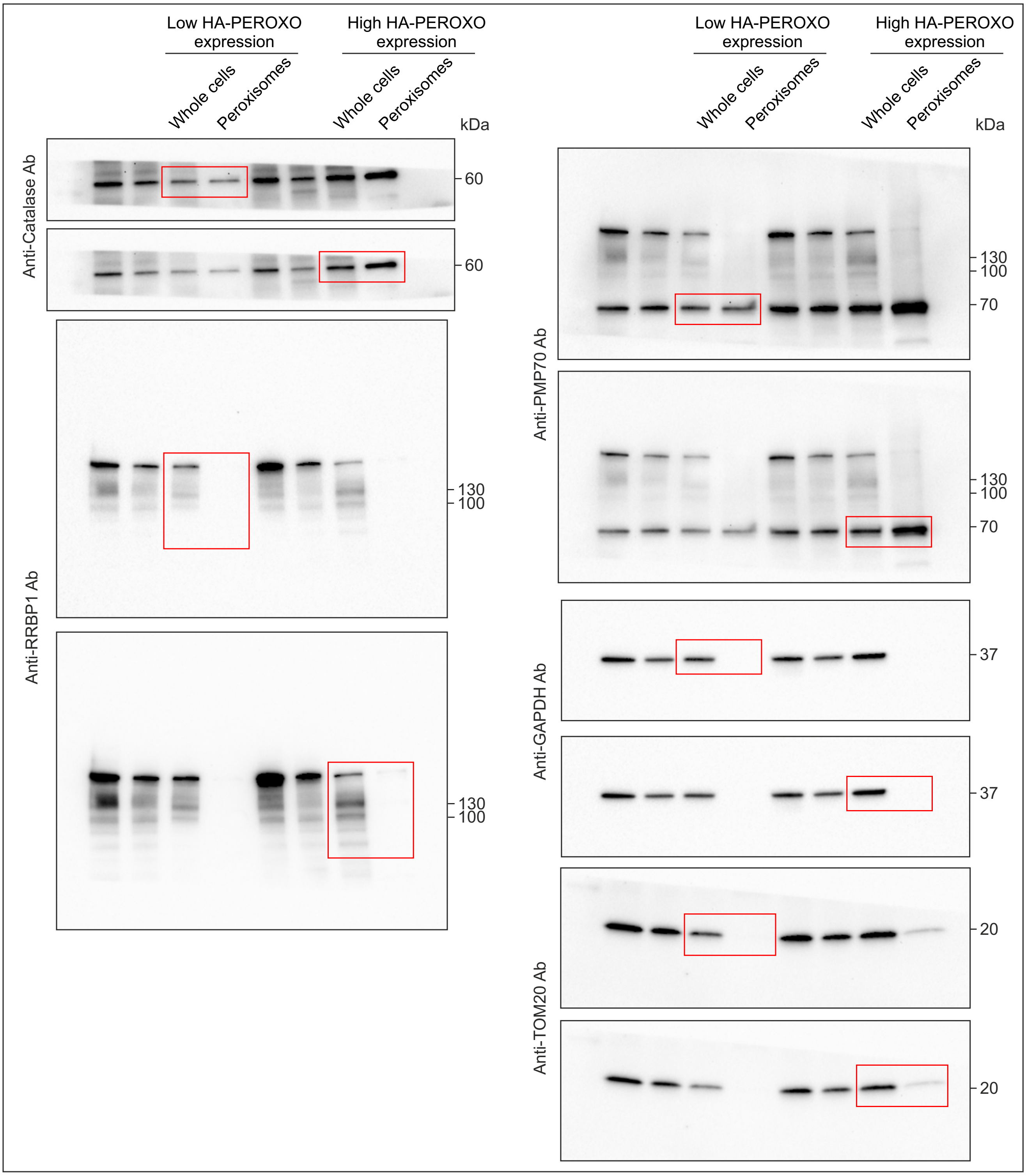

Blot 2: 10% SDS PAGE. The blot was cut onto three parts, PEX3 was detected on the middle part of the blot.
Calnexin was detected on the upper part of the blot. PEX16 was detected on the lowere part of the blot.

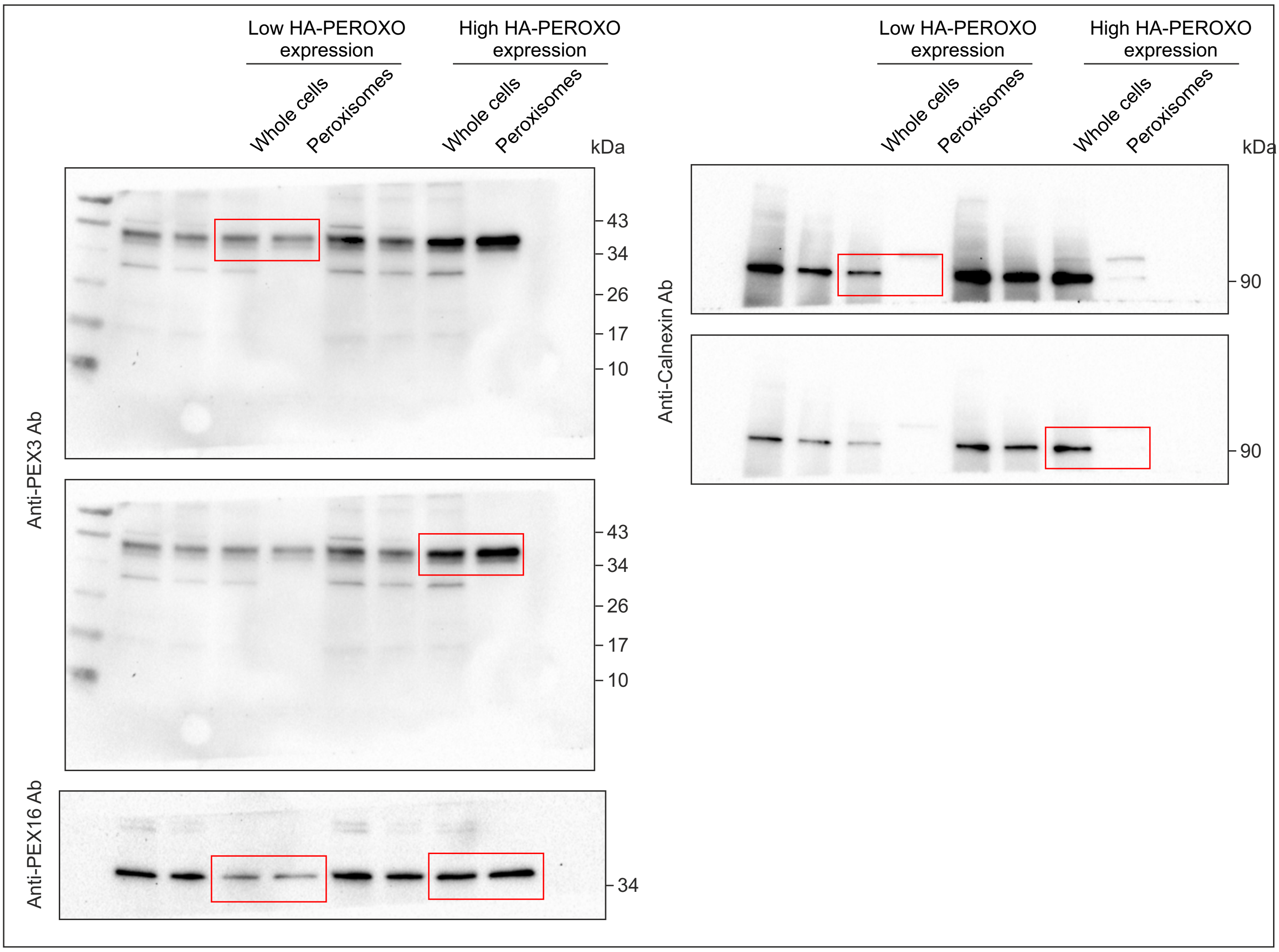

### Unprocessed western blot images related to Figure 6C

Western blot analysis of RRBP1 silencing in HEK293T cells stably expressing GFP-APEX2-ePTS1.
The cells were transfected with corresponding esiRNAs for 24 or 48 h.
Luciferase (Luc) esiRNA was used as a negative control.
Areas presented in the figure are indicated.

10% SDS PAGE. The membrane was cut onto two parts. The upper part was used to detect RRBP1 and the lower part was used for β-tubulin detection..

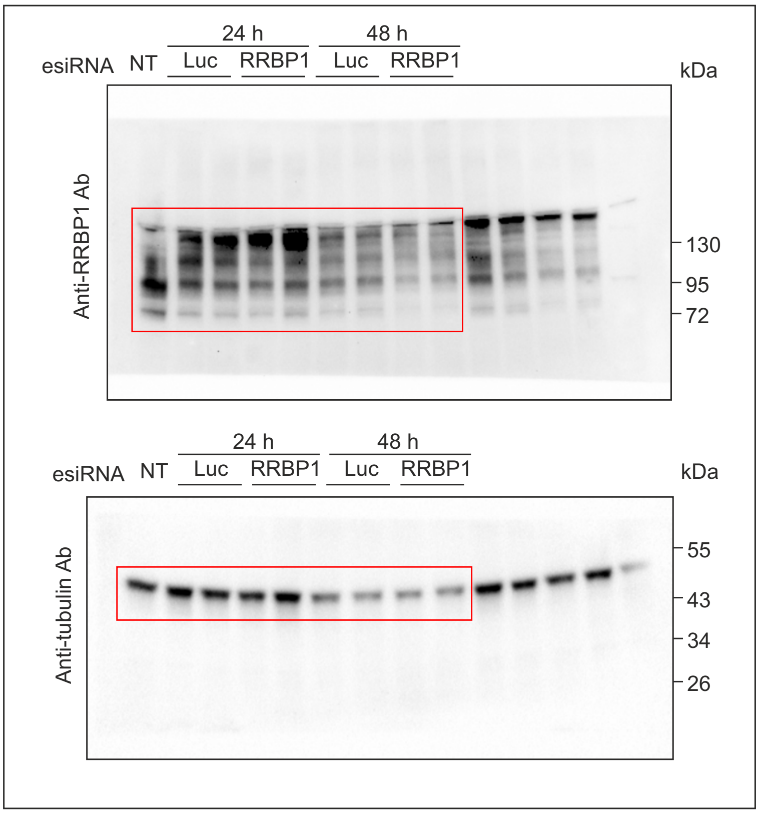

### Unprocessed western blot images related to Figure 7A

Western blot analysis of RRBP1 KO or parental HEK293T cells stability expressing GFP-APEX2-ePTS1 construct before and after treatment with 100 μM chloroquine for 3 h.
Areas presented in the figure are indicated.

Blot 1: 10 % SDS PAGE. RRBP1, PMP70 and GAPDH were detected on the same blot in the folllowing order: RRBP1, PMP70 and GAPDH last.

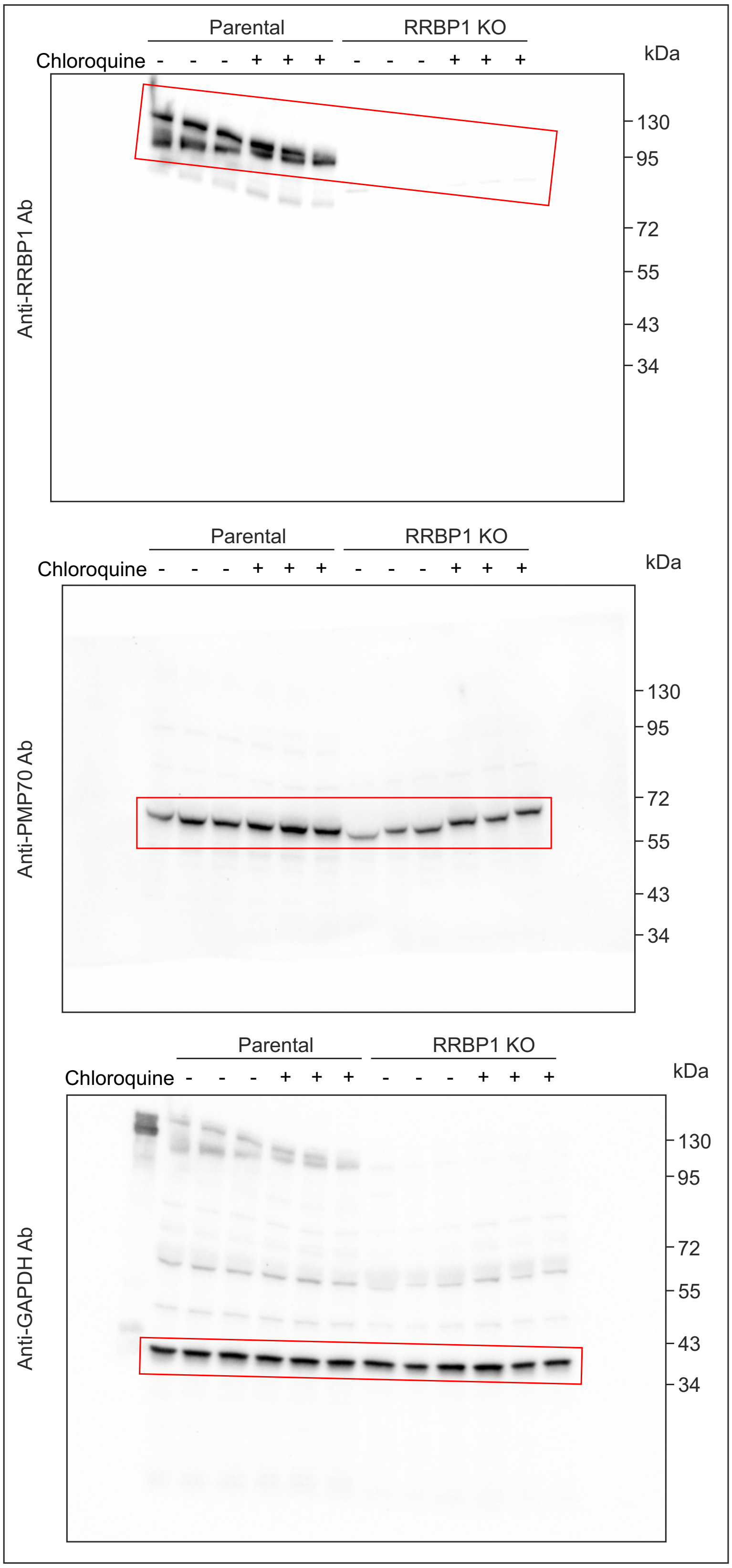

Blot 2: 10% SDS PAGE. The membrane was cut onto two parts. LC3B was detected from the bottom part.
ATG5, PEX3 and GAPDH were detected on the upper part in the folllowing order: PEX3, GAPDH and ATG5 last.

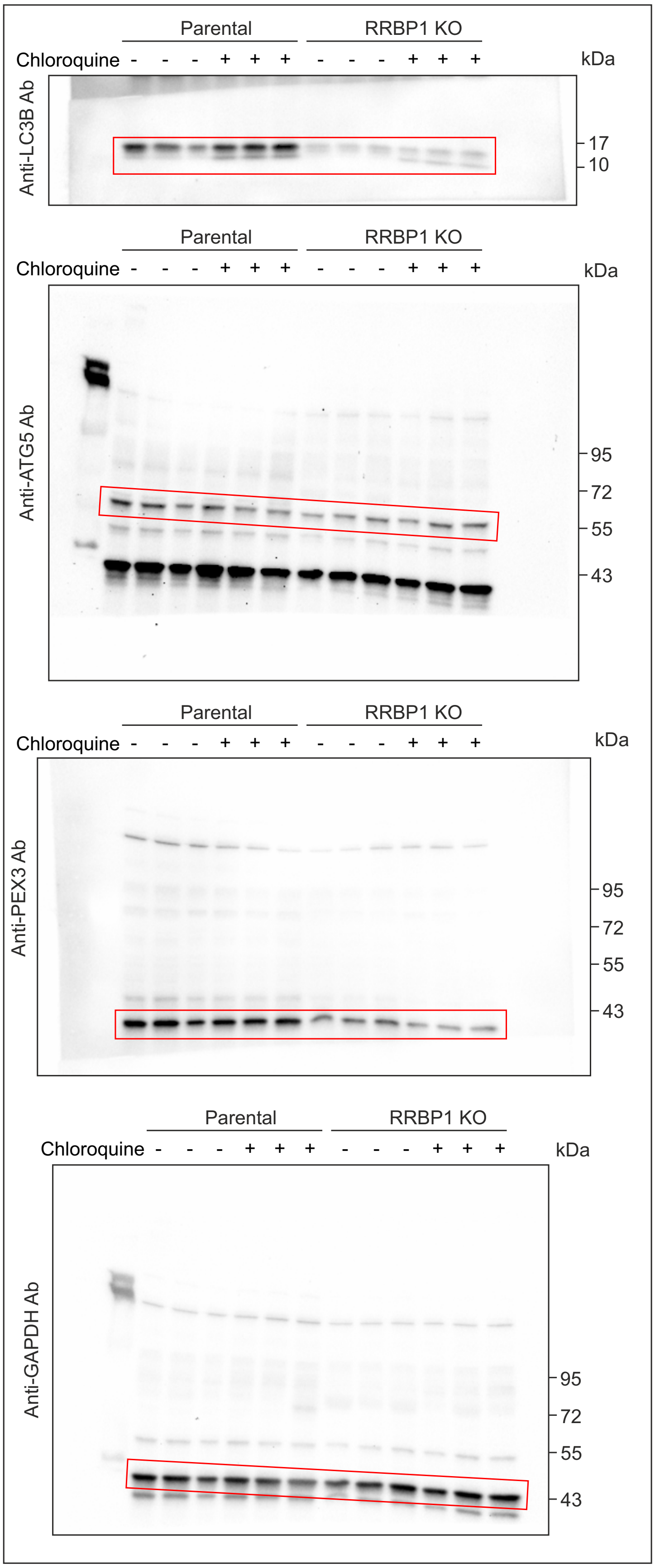

#### Unprocessed western blot images related to Figure 7C

Western blot analysis of RRBP1 KO or parental HEK293T cells stability expressing GFP-APEX2-ePTS1 construct before and after treatment with 10 µM MG132 for 4 h.
Areas presented in the figure are indicated.

10 % SDS PAGE. Stain-Free total protein staining was used as a loading control. The membrane was cut onto two parts. Upper part was used to detect RRBP1. Lower part was used for the detection of PEX3. Then the lower part was cut onto two parts and PEX16 was detected.

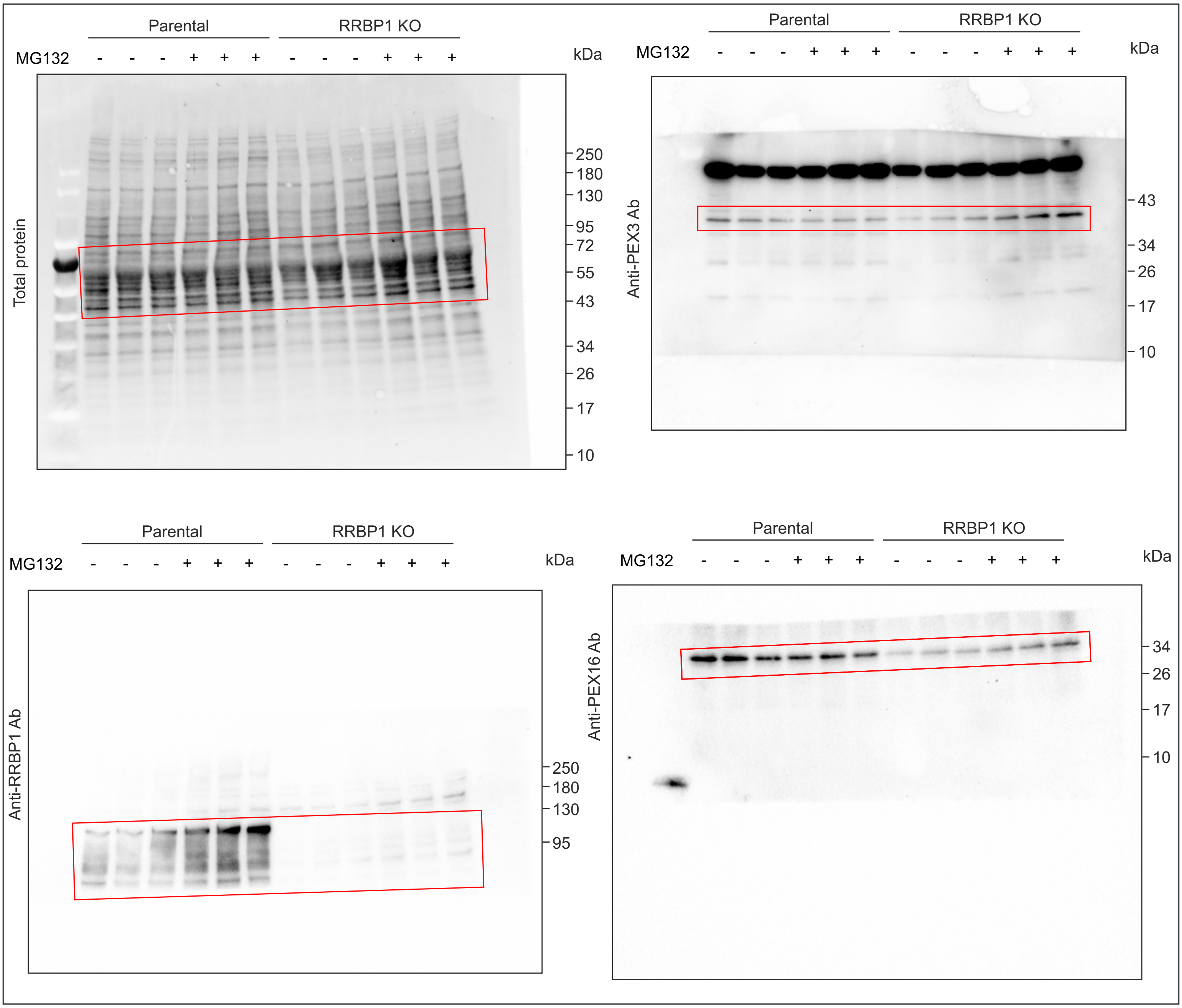

#### Unprocessed western blot images related to Supplementary Figure 1A

Western blotting analysis of HEK293T single cell clones stably expressing GFP-APEX2-ePTS1 or GFP-APEX2 constructs.
Areas presented in the figure are indicated.

Blot 1: 10% SDS PAGE. PEX19, GFP and GAPDH were detected on the same blot in the folllowing order: PEX19, GAPDH and GFP last.
Blot 2: 10% SDS PAGE. PEX3 and tubulin were detected on the same blot in the folllowing order: PEX3 first and then tubulin.

**Blot 1**

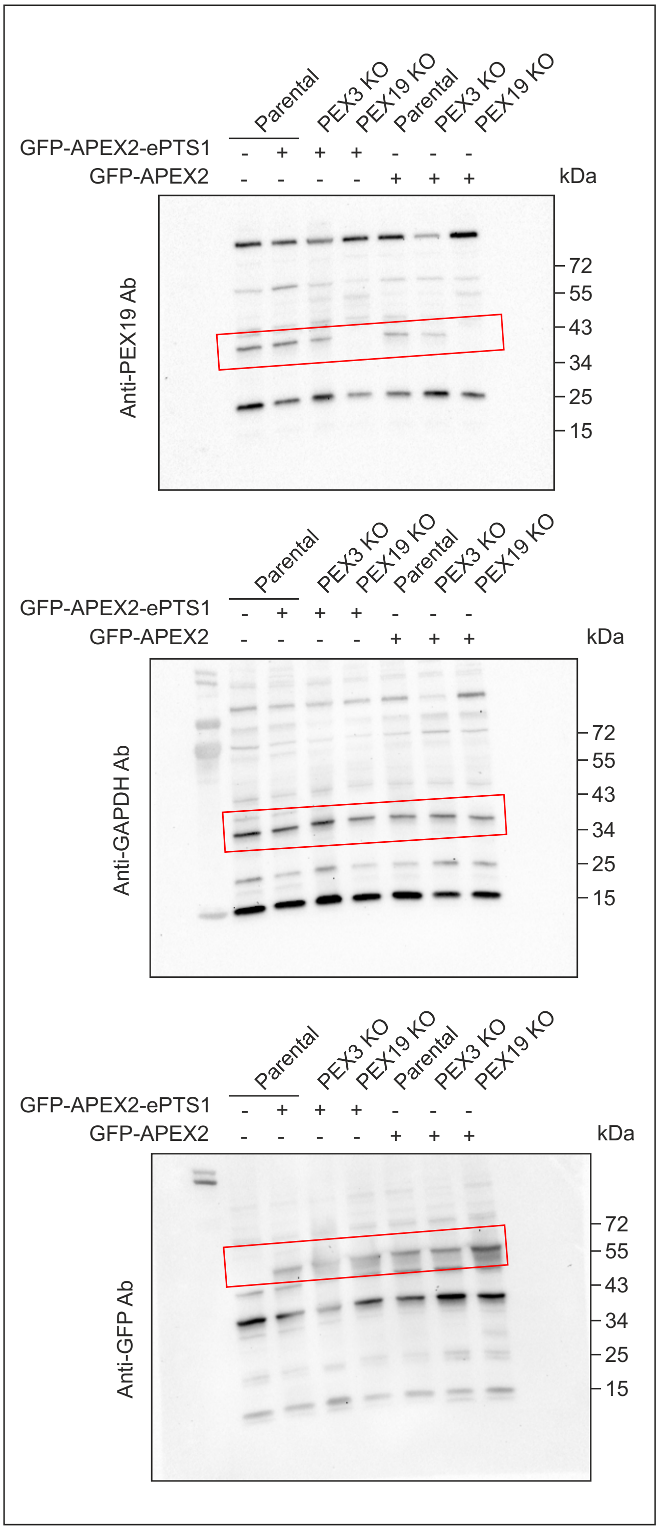


**Blot 2**

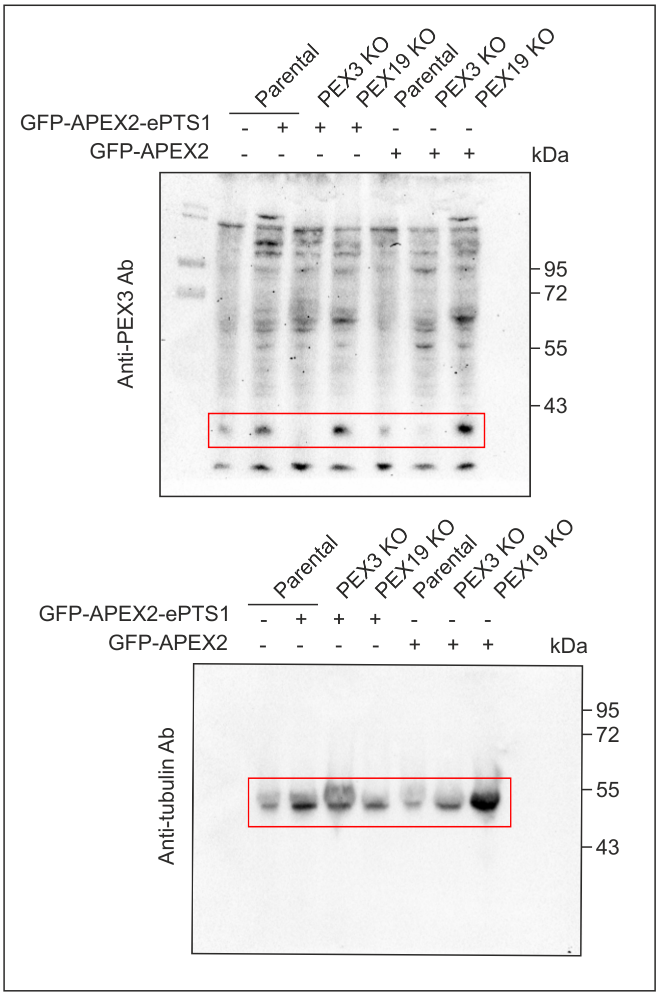

Blot 3: 10% SDS PAGE. PEX5 and GAPDH were detected on the same blot in the folllowing order: PEX5 first and then GAPDH.

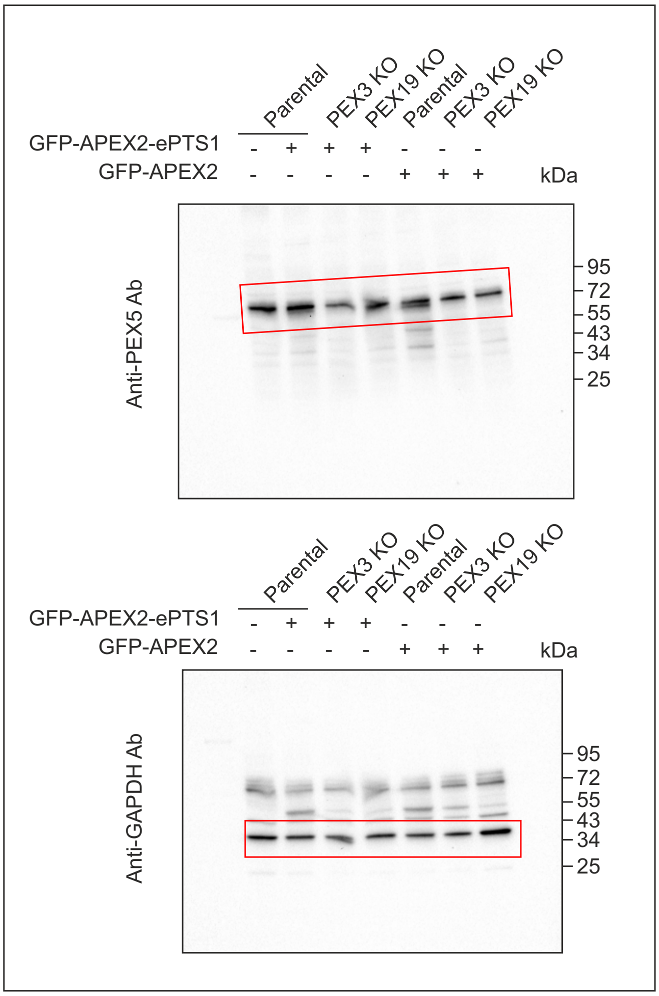

### Unprocessed western blot images related to Supplementary Figure 1E

Western blot analysis of GFP-APEX2-ePTS protein level in HEK293T cells stably expressing GFP-APEX2-ePTS1 before and after 13 days of transfection with Cas9-mCherry plasmid and gRNA targeting PEX19, PEX5 or RRBP1. Anti-GFP antibody was used to detect the GFP-APEX2-ePTS protein.
Areas presented in the figure are indicated.

Blot 1: 10% SDS PAGE. GFP and GAPDH were detected on the same blot.
Blot 2: 10% SDS PAGE. GFP and GAPDH were detected on the same blot.

**Blot 1**

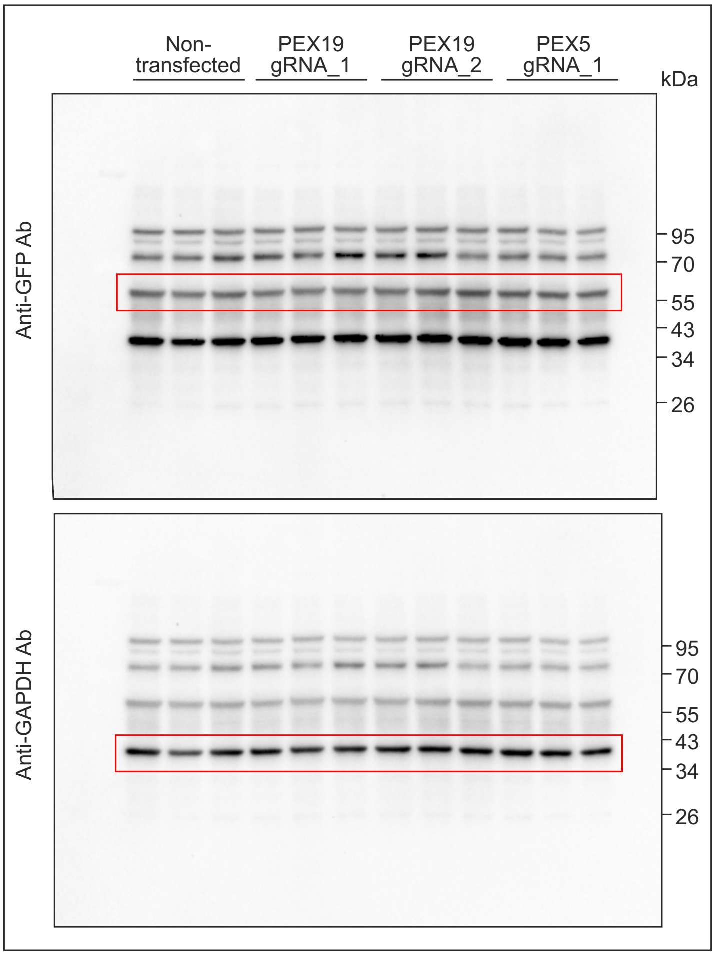


**Blot 2**

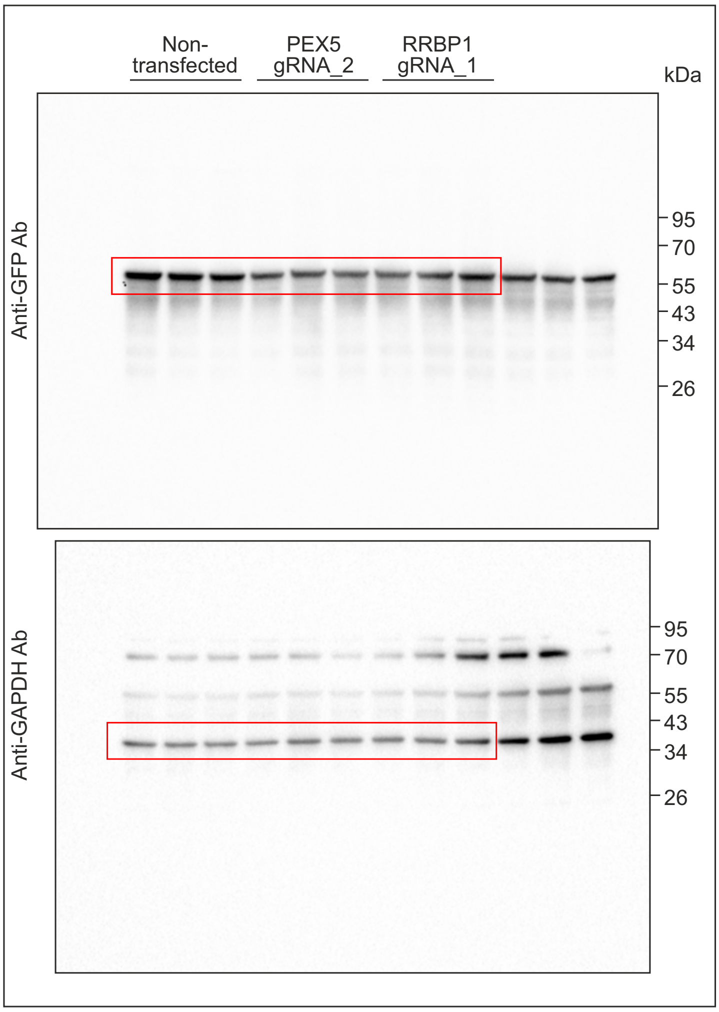

#### Unprocessed western blot images related to Supplementary Figure 2B

Western blot analysis of RRBP1 KO or parental HEK293T cells stably expressing GFP-APEX2-ePTS1 before or after puromycin treatment (10 µg/ml, 20 min).
Areas presented in the figure are indicated.

10% SDS PAGE. Puomycin and GAPDH were detected on the same blot.

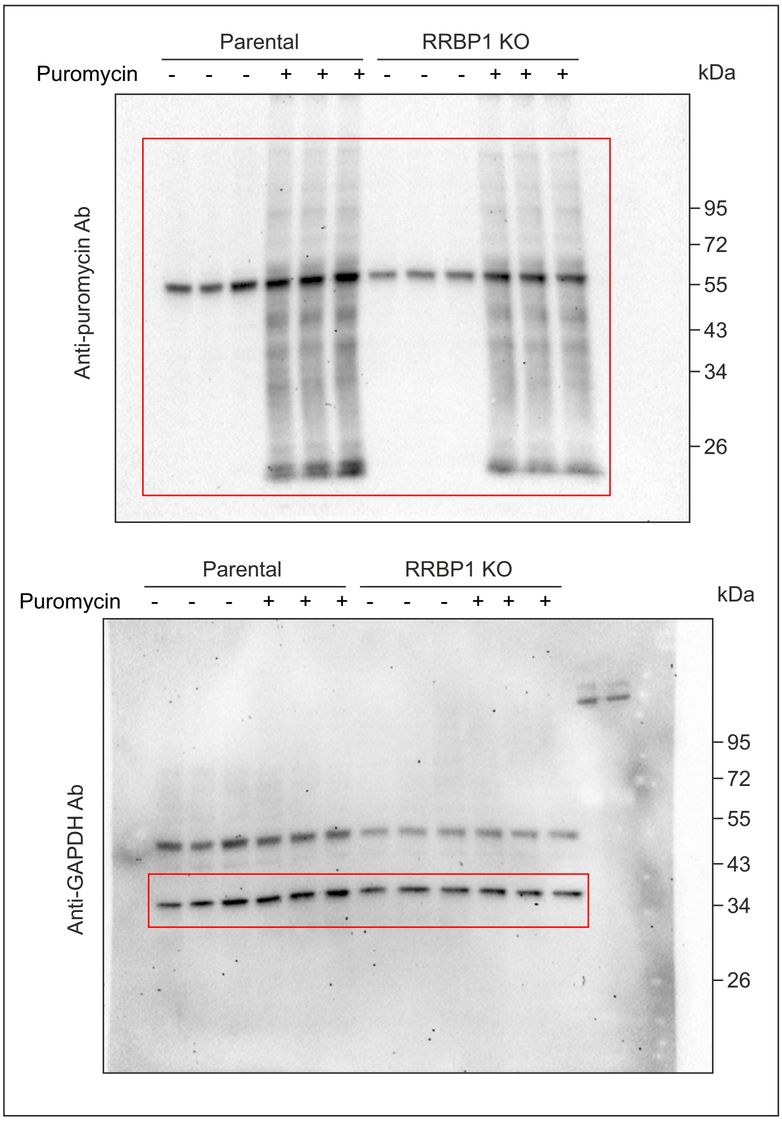

### Unprocessed western blot images related to Supplementary Figure 2D

Western blot analysis of RRBP1 KO HEK293T cells stably expressing GFP-APEX2-ePTS1.
Parental, PEX3 and PEX19 KO HEK293T cells not expressing peroxisomal targeted construct were used as controls. GAPDH was used as a loading control.
Areas presented in the figure are indicated.

Blot 1: 10% SDS PAGE. SEC61, IRE1, PXMP2 and GAPDH were detected on the same blot.

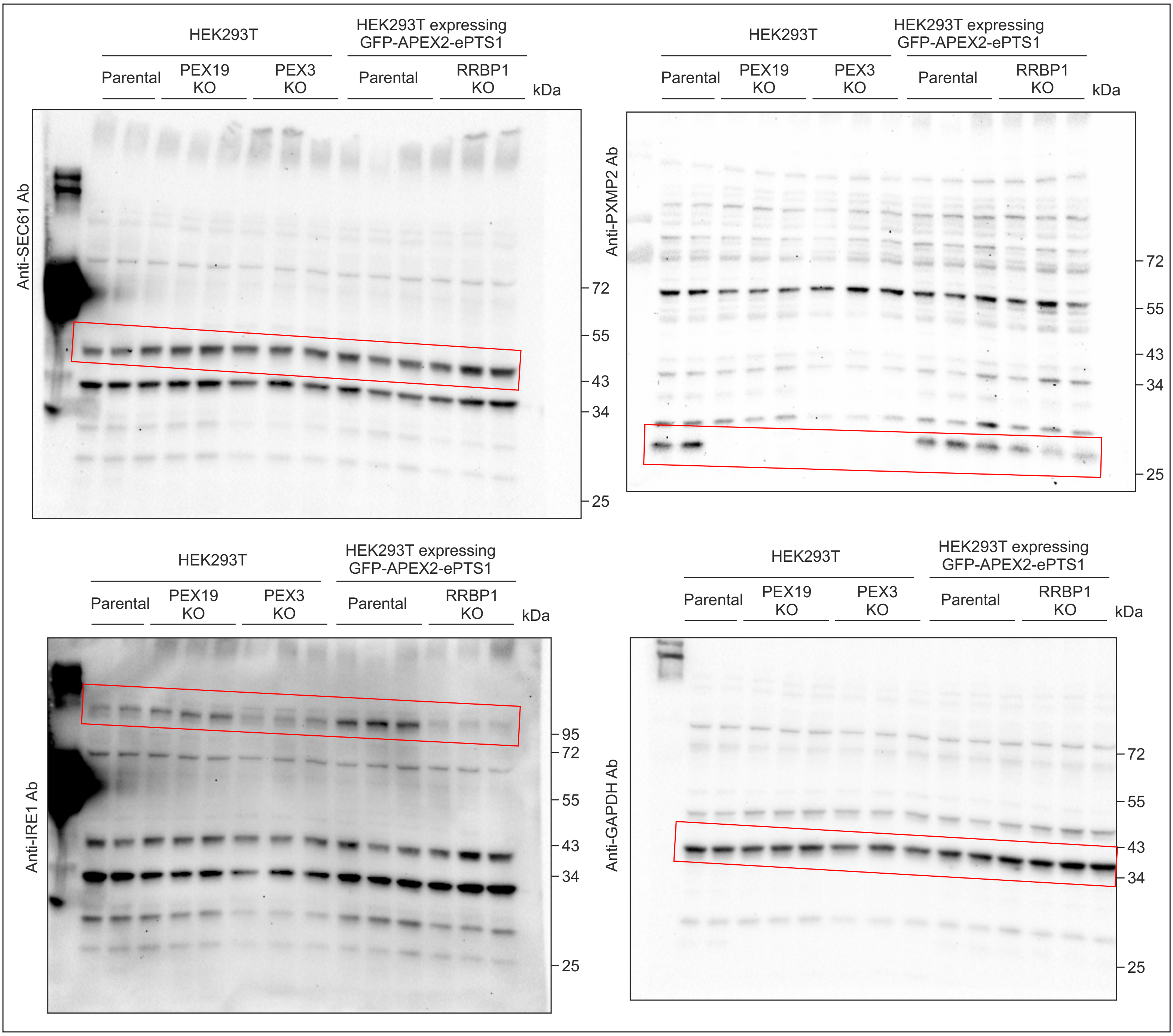

Blot 2: 10% SDS PAGE. BiP, RPL7 and GAPDH were detected on the same blot.

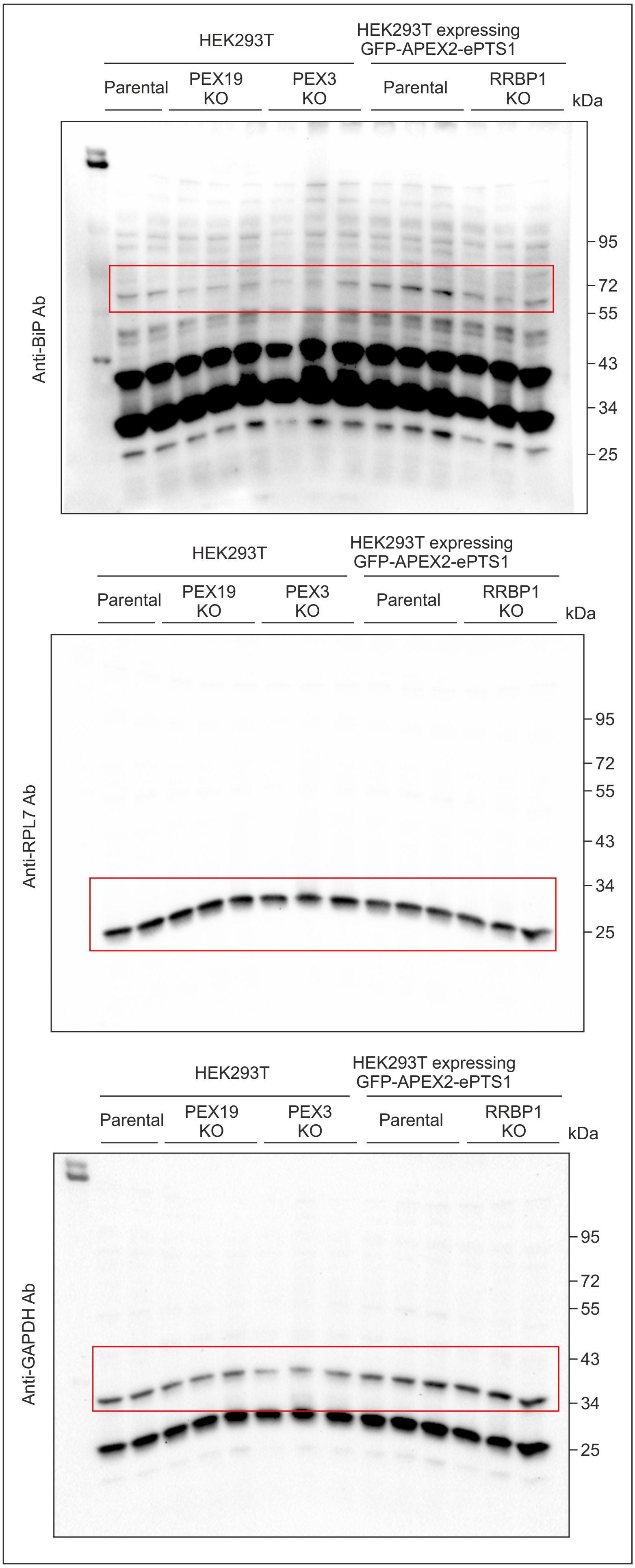
