## Supplementary material for "Ribosome-binding protein 1, RRBP1, maintains peroxisome biogenesis": Blot Transparency File

### Unprocessed western blot images related to Figure 2A

Western blot analysis of RRBP1 KO HEK293T cells stably expressing GFP-APEX2-ePTS1.
Parental, PEX3 and PEX19 KO HEK293 cells without peroxisomal targeted construct were used as controls.
Areas presented in the figure are indicated.

Blot 1: 7.5% SDS PAGE. RRBP1 and GAPDH were detected on the same blot in the folllowing order: RRBP1 first and GFP last.

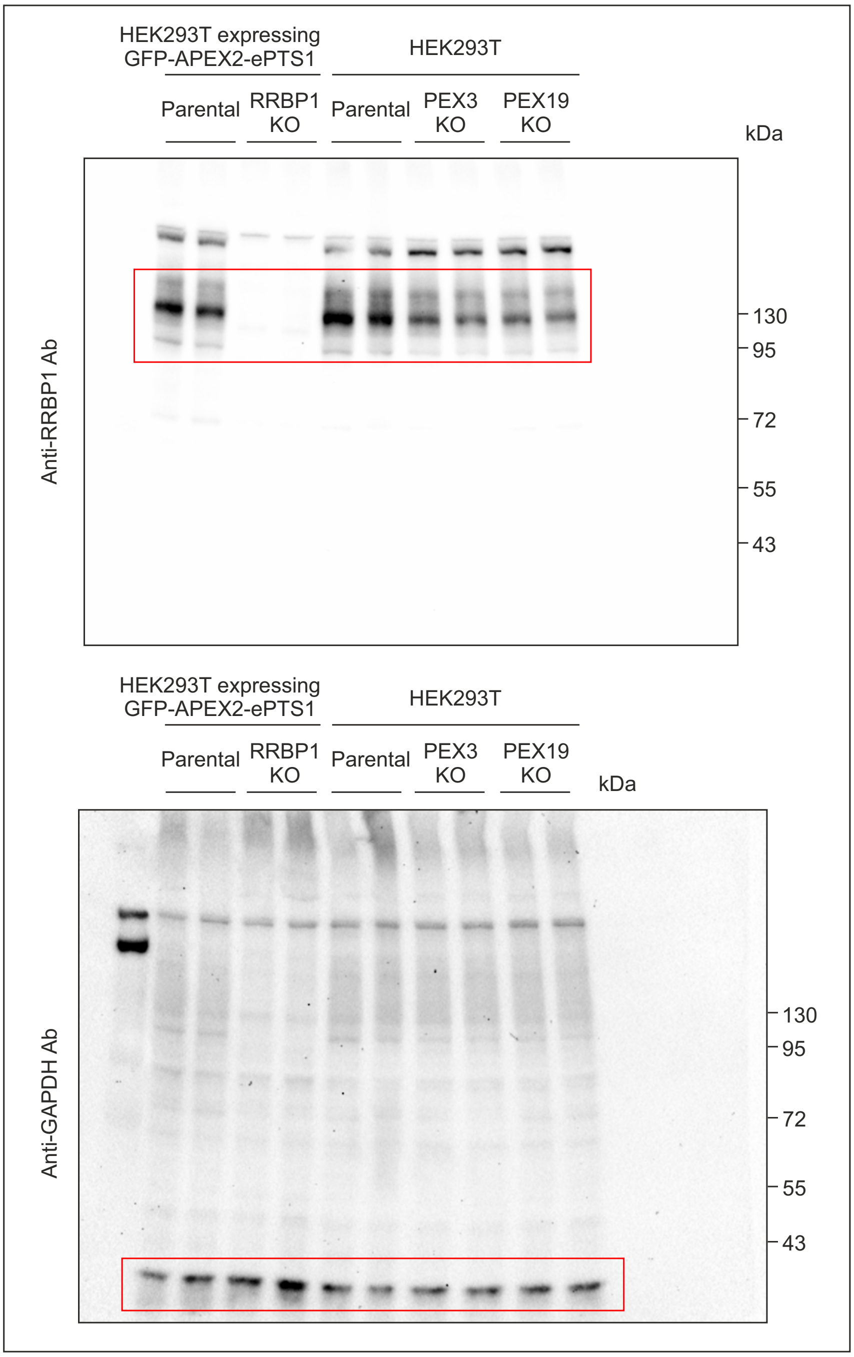

Blot 2: 10% SDS PAGE. PEX5, catalase, TOM20 and GAPDH were detected on the same blot in the folllowing order: catalase, PEX5, TOM20 and GAPDH last.

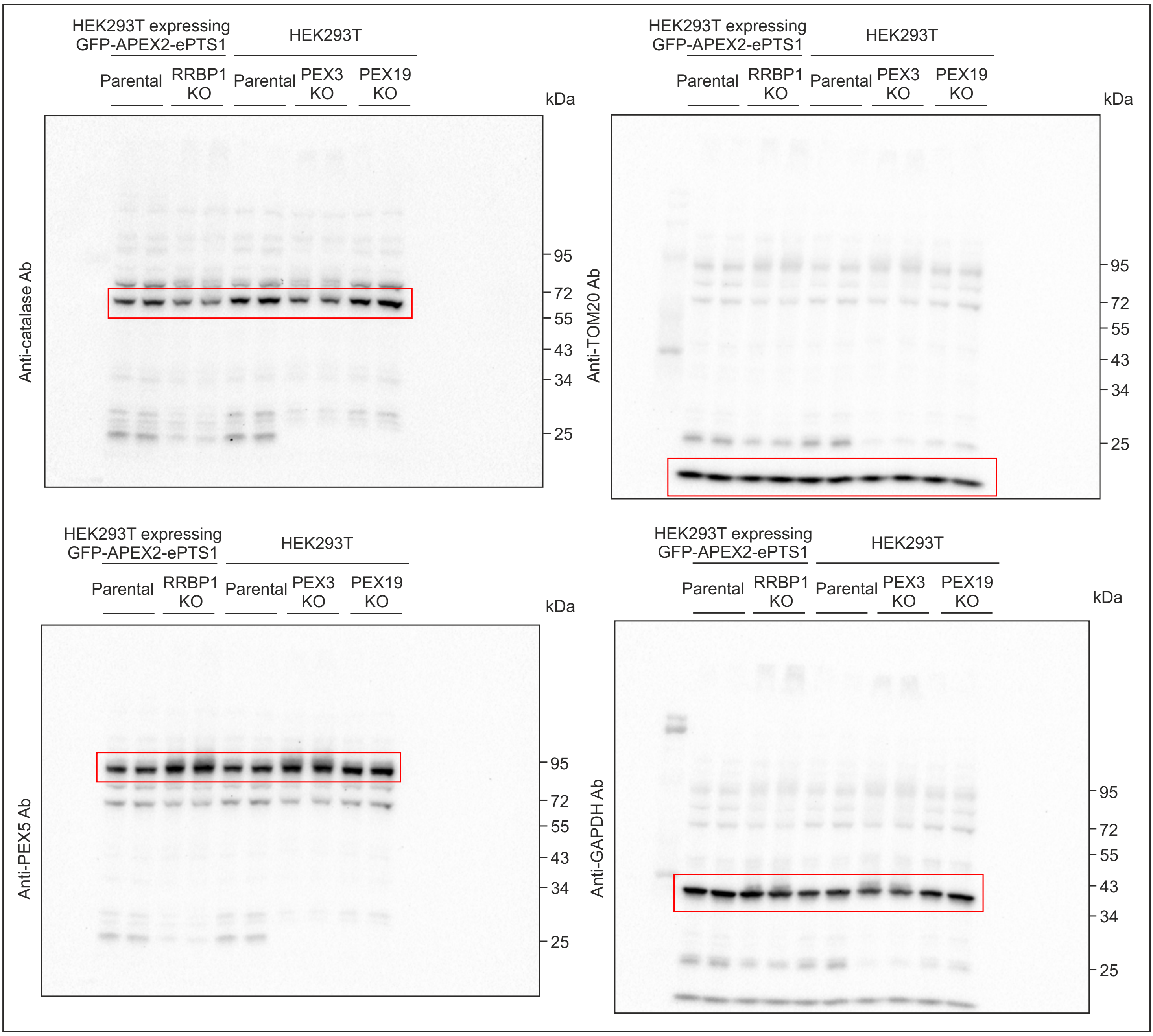

Blot 3: 10% SDS PAGE. ACOX1, PEX14 and GAPDH were detected on the same blot in the folllowing order: ACOX1 first, then GAPDH and PEX14 last.

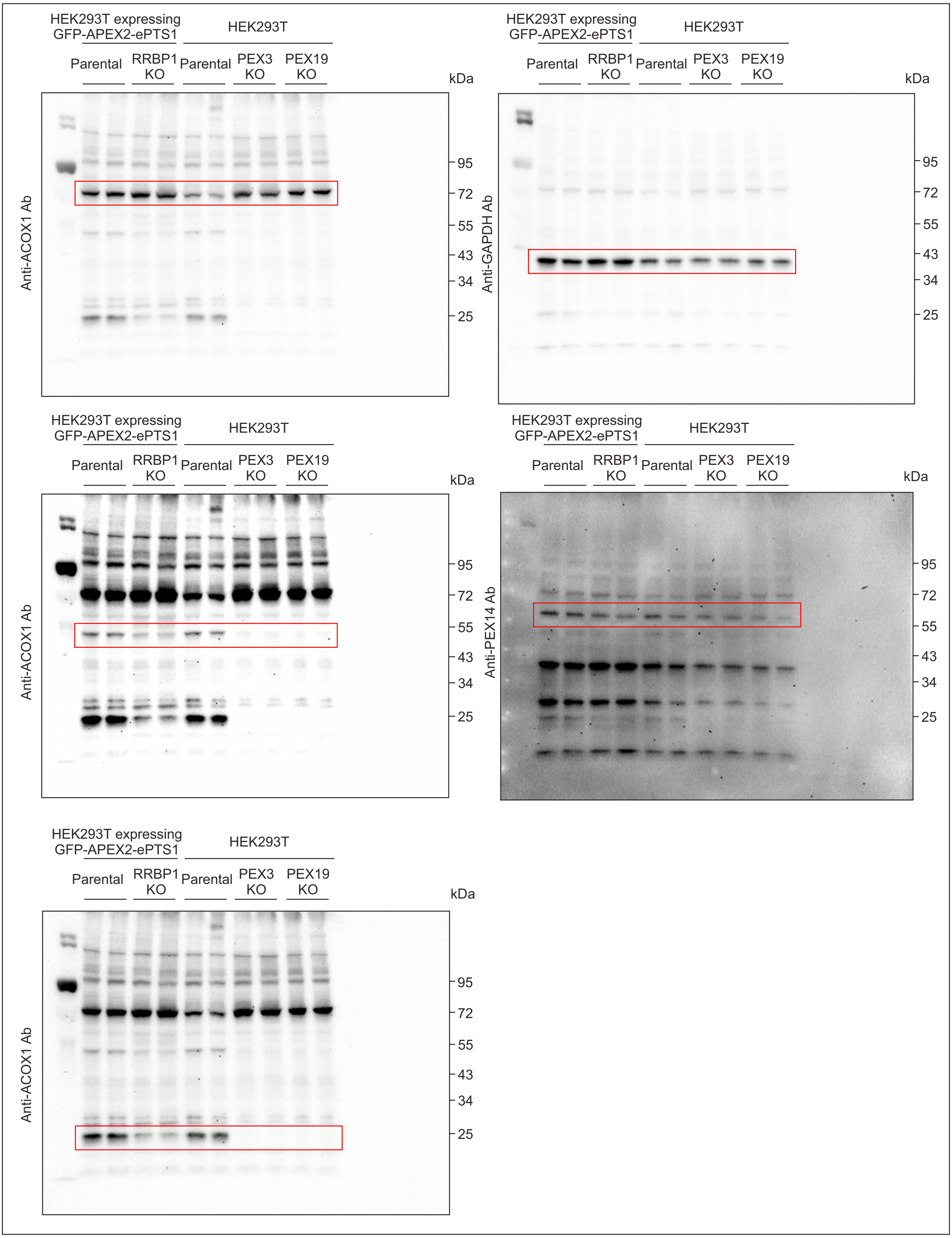

Blot 4: 10% SDS PAGE. PEX3, calnexin, VAPB and GAPDH were detected on the same blot in the folllowing order: calnexin, PEX3, GAPDH and VAPB last.

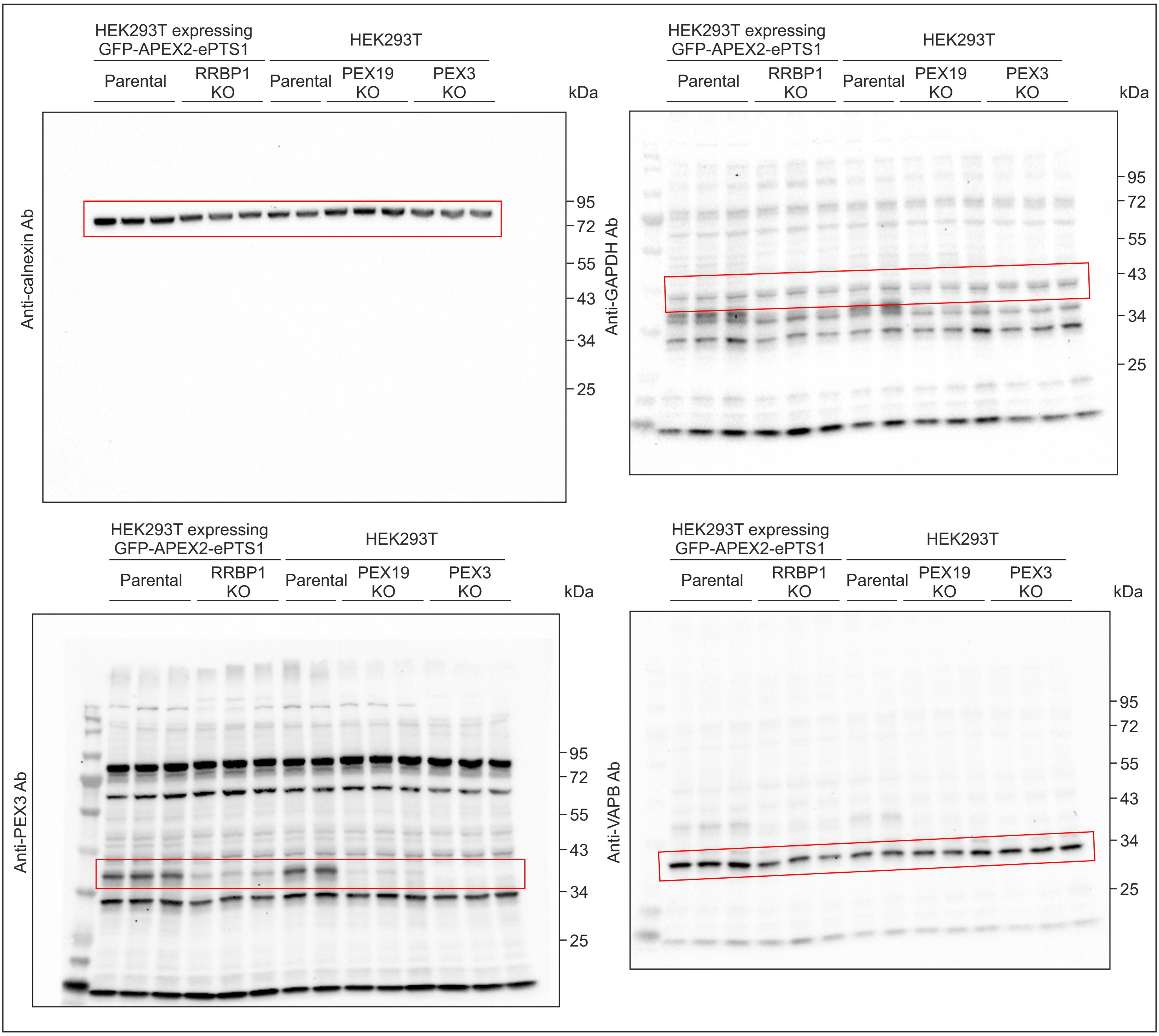

Blot 5: 10% SDS PAGE. PEX19 and GAPDH were detected on the same blot in the folllowing order: PEX19 first and then GAPDH.

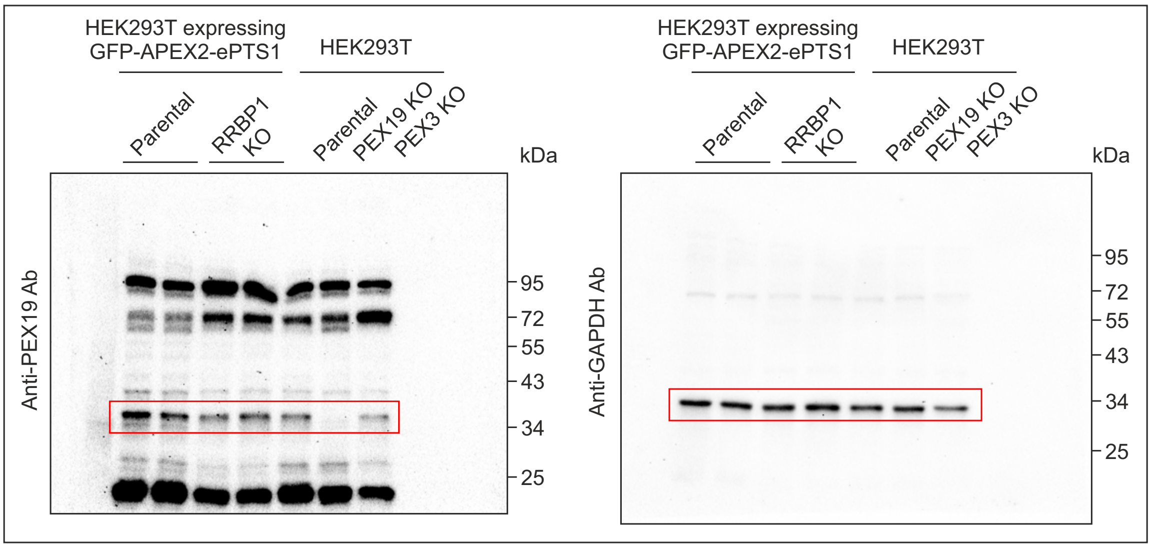


Blot 6: 7.5% SDS PAGE. ACAA1 and GAPDH were detected on the same blot in the folllowing order: ACAA1 first and then GAPDH.

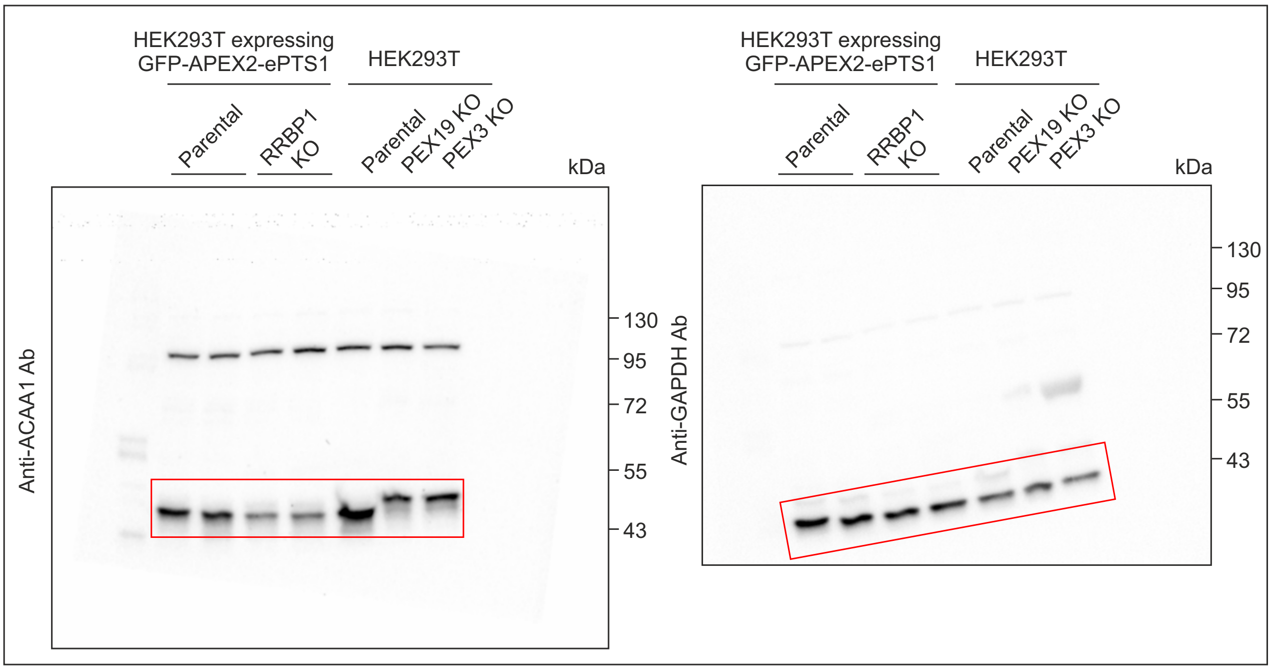

### Unprocessed western blot images related to Figure 4A

Western blot analysis of RRBP1 silencing in HEK293T cells stably expressing GFP-APEX2-ePTS1.
The cells were transfected with corresponding esiRNAs for 72 h. GFP esiRNA was used as a negative control, esiRNA PEX5 was used as a positive control.
Areas presented in the figure are indicated.

Blot 1: 7.5% SDS PAGE. GFP, RRBP1, PEX5, ACAA1 and GAPDH were detected on the same blot in the folllowing order: RRBP1, ACAA1, PEX4, GFP and GAPDH last.

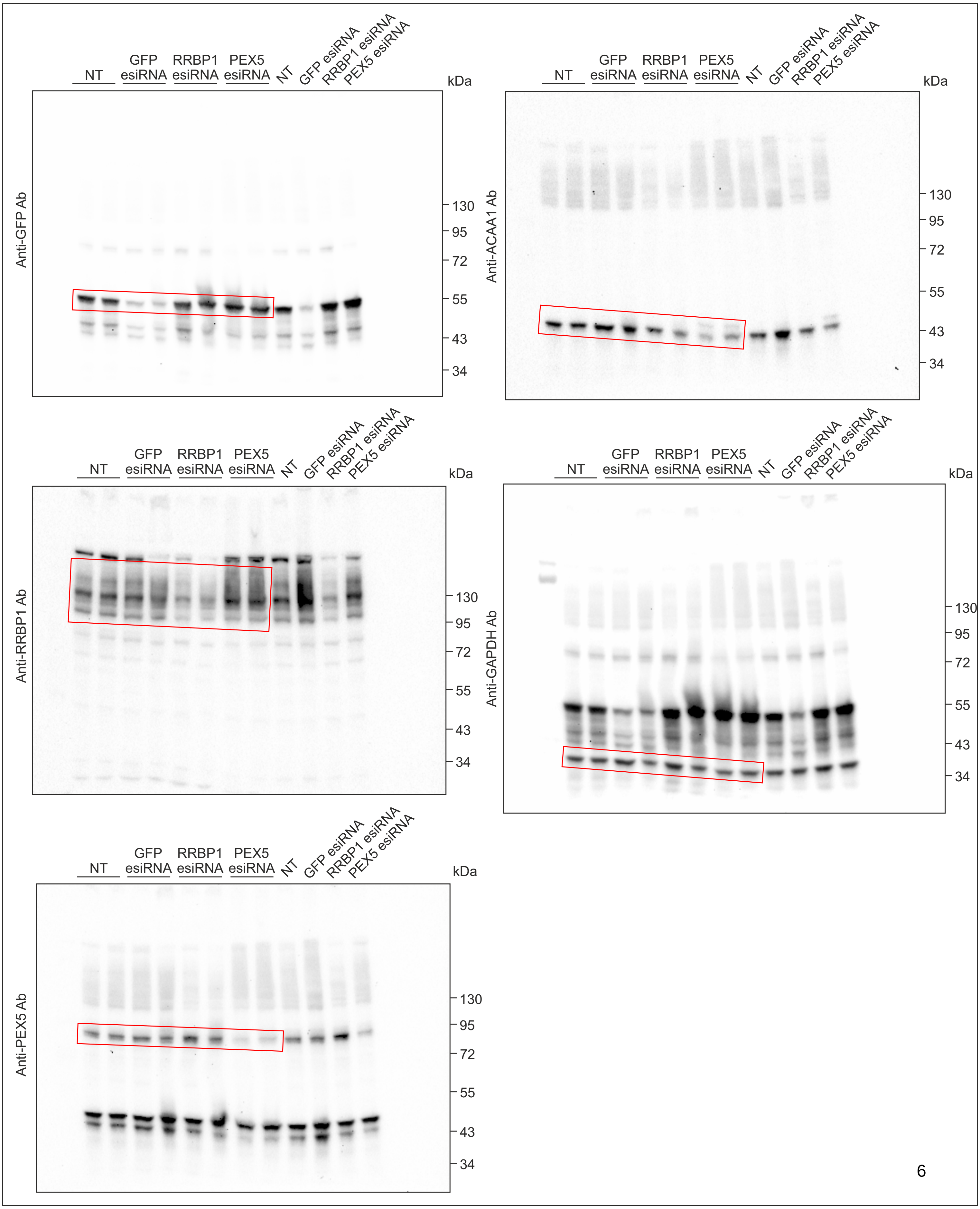

Blot 2: 10% SDS PAGE. ACOX1, catalase and tubulin were detected on the same blot in the folllowing order: ACOX1 first, then catalase and tubulin last.

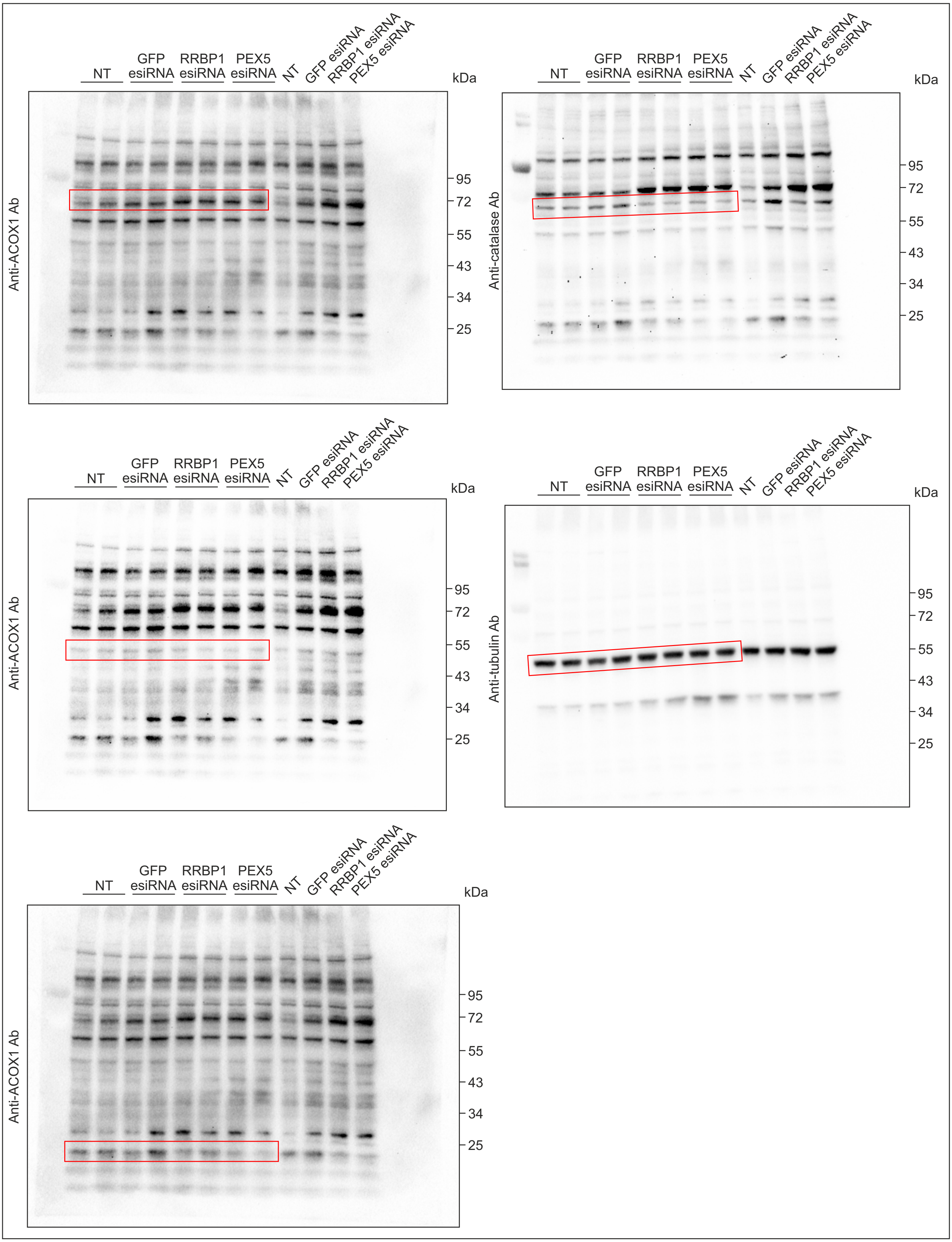

#### Unprocessed western blot images related to Figure 5C

Western blot of HEK293T cells expressing 3×Myc-EGFP-PEX26 (HA-PEROXO) and immunopurified peroxisomes.
Areas presented in the figure are indicated.

Blot 1: 10% SDS PAGE. The blot was cut into four parts. PMP70 and RRBP1 were detected on the upper part of the blot in the folllowing order: RRBP1 first and then PMP70. Catalase was detected on the second part of the blot. GAPDH was detected on the third part of the blot. TOM20 was detected on the lowest part of the blot.

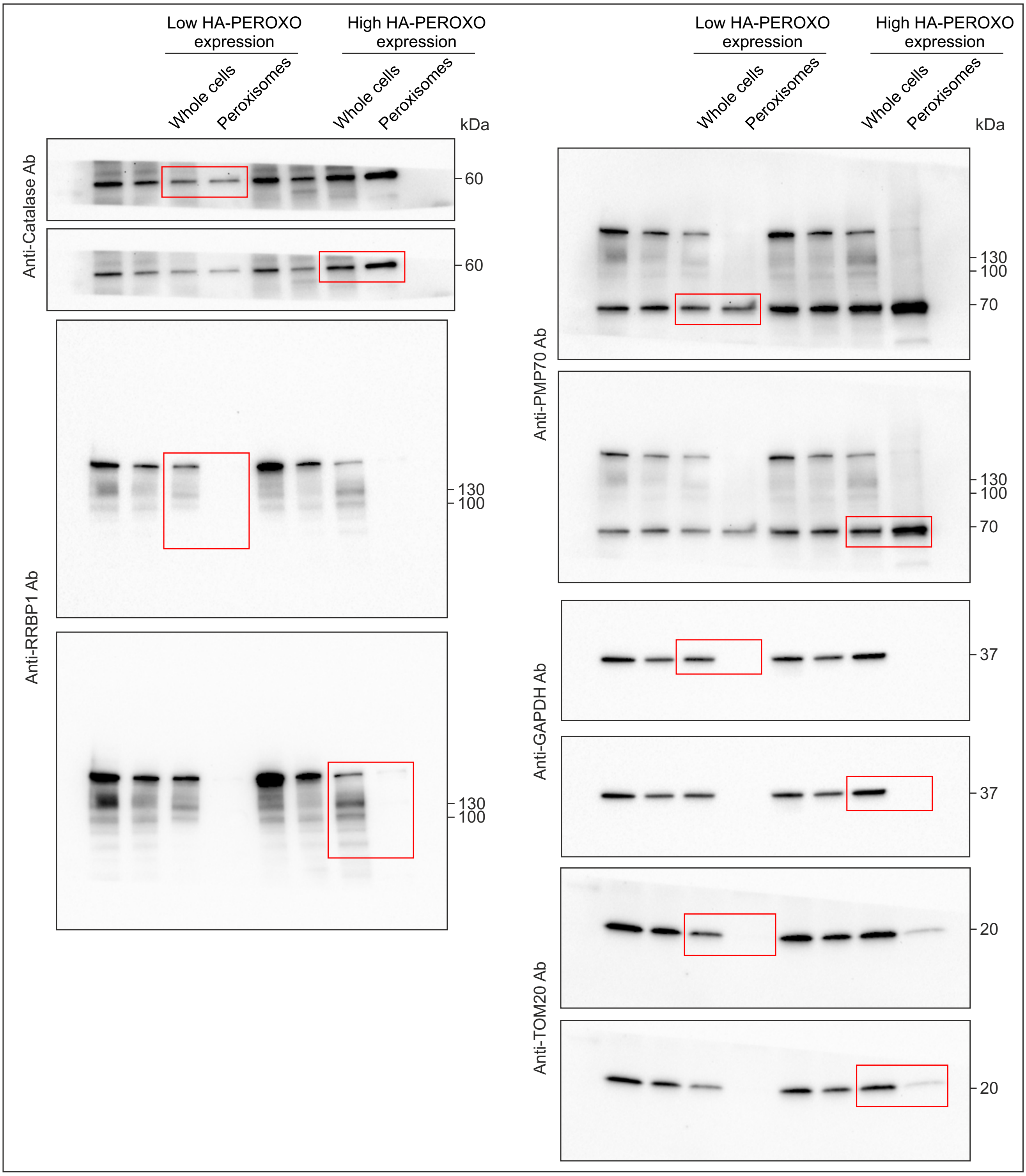

Blot 2: 10% SDS PAGE. The blot was cut onto three parts, PEX3 was detected on the middle part of the blot.
Calnexin was detected on the upper part of the blot. PEX16 was detected on the lowere part of the blot.

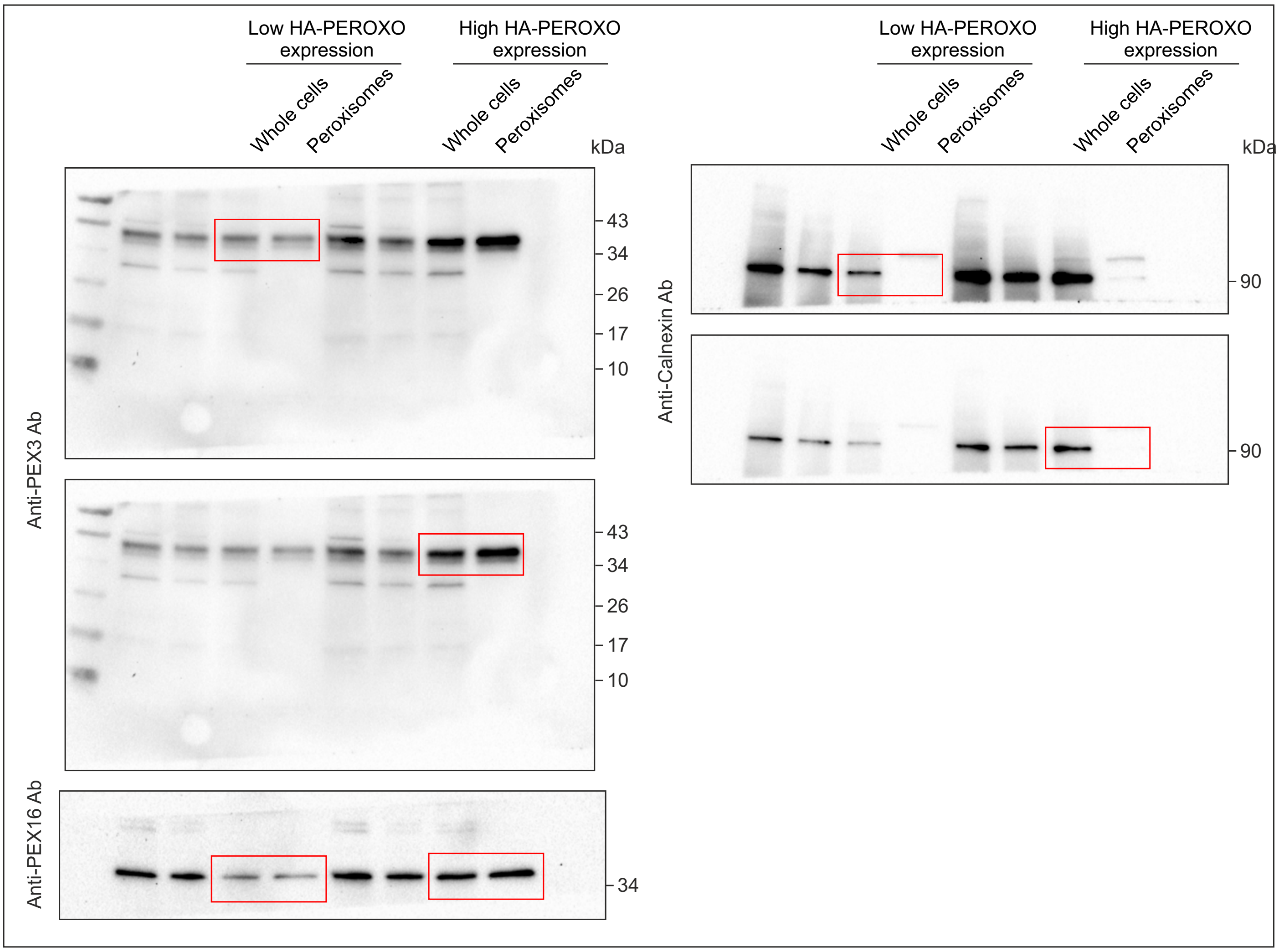

### Unprocessed western blot images related to Figure 6C

Western blot analysis of RRBP1 silencing in HEK293T cells stably expressing GFP-APEX2-ePTS1.
The cells were transfected with corresponding esiRNAs for 24 or 48 h.
Luciferase (Luc) esiRNA was used as a negative control.
Areas presented in the figure are indicated.

10% SDS PAGE. The membrane was cut onto two parts. The upper part was used to detect RRBP1 and the lower part was used for β-tubulin detection..

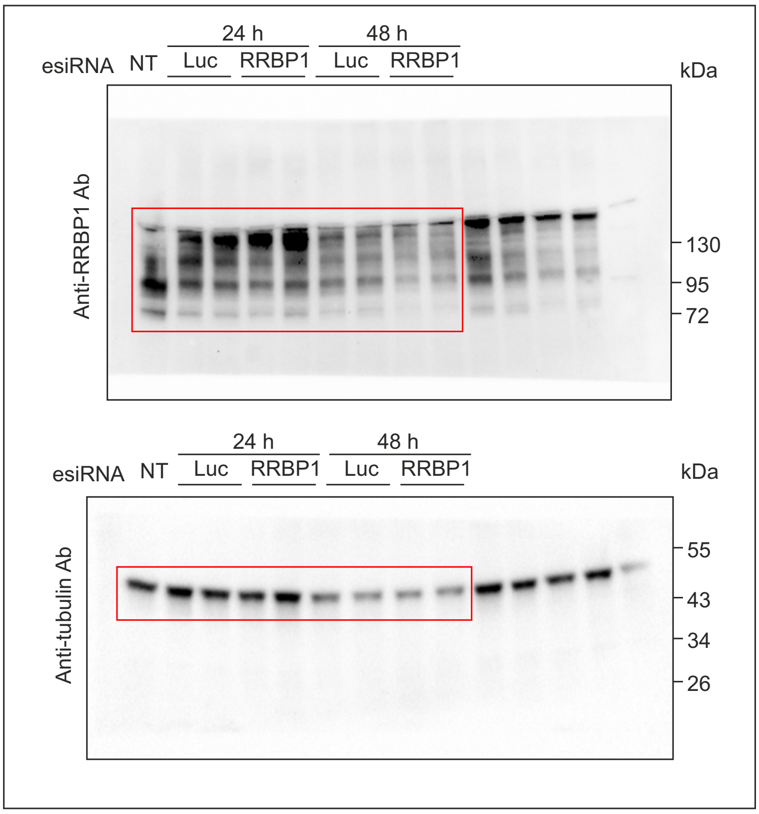

### Unprocessed western blot images related to Figure 7A

Western blot analysis of RRBP1 KO or parental HEK293T cells stability expressing GFP-APEX2-ePTS1 construct before and after treatment with 100 μM chloroquine for 3 h.
Areas presented in the figure are indicated.

Blot 1: 10 % SDS PAGE. RRBP1, PMP70 and GAPDH were detected on the same blot in the folllowing order: RRBP1, PMP70 and GAPDH last.

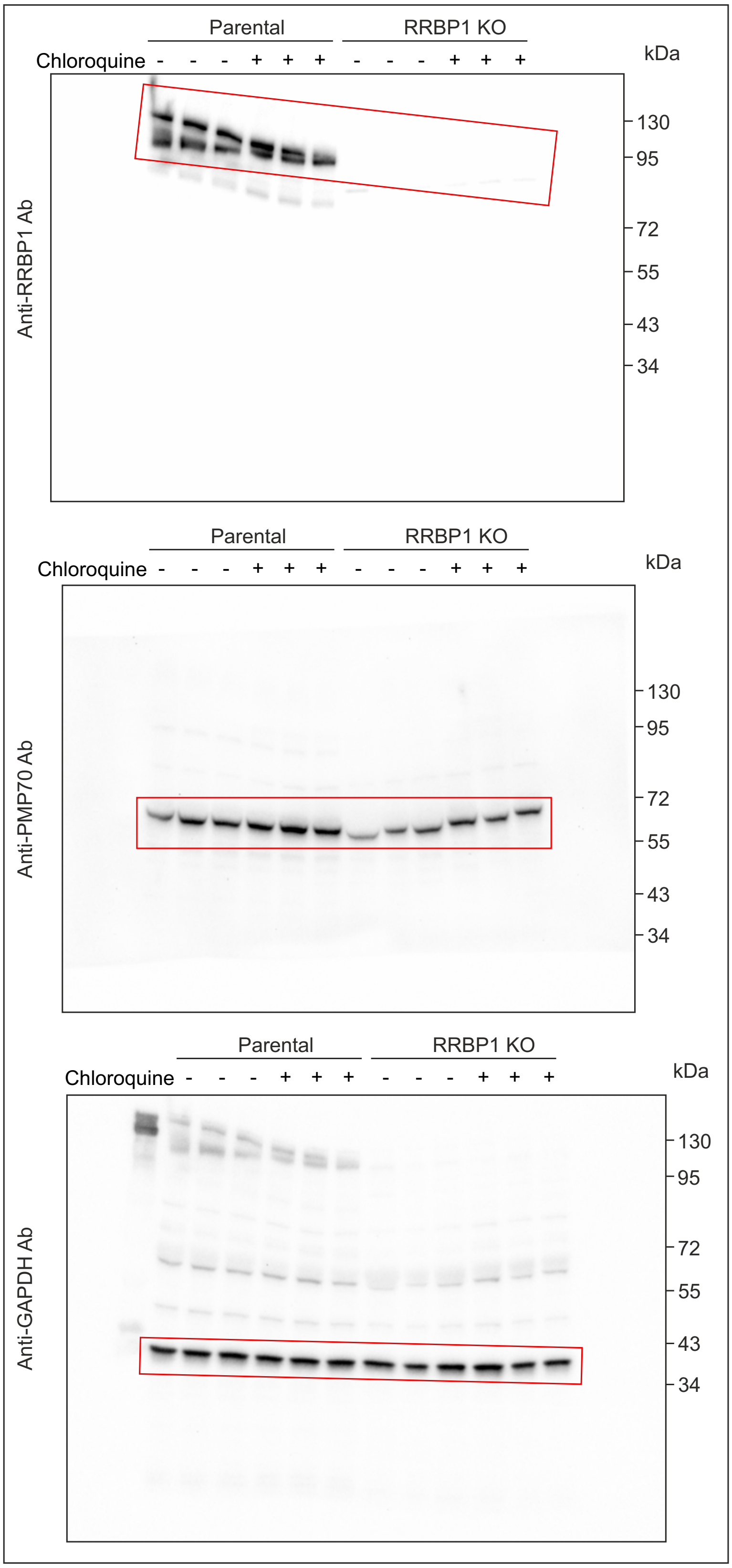

Blot 2: 10% SDS PAGE. The membrane was cut onto two parts. LC3B was detected from the bottom part.
ATG5, PEX3 and GAPDH were detected on the upper part in the folllowing order: PEX3, GAPDH and ATG5 last.

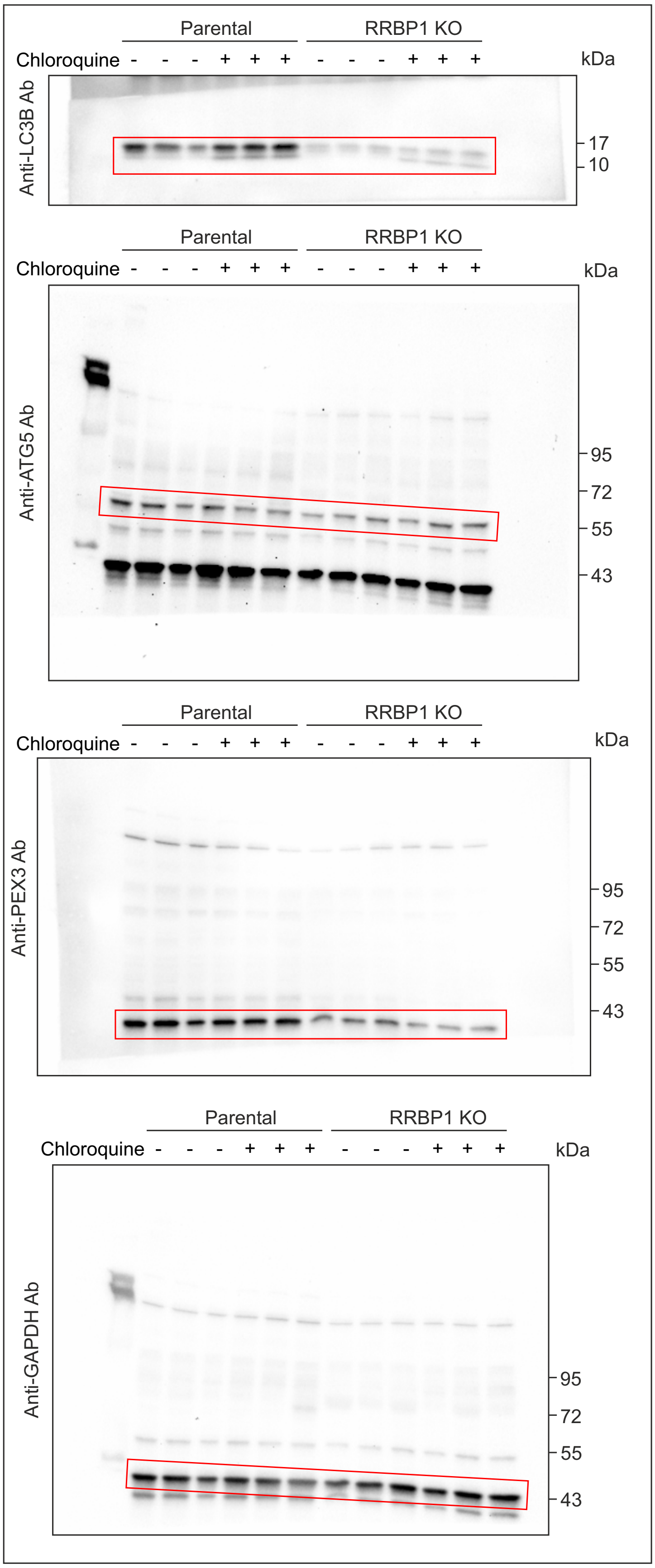

### Unprocessed western blot images related to Figure 7C

Western blot analysis of RRBP1 KO or parental HEK293T cells stability expressing GFP-APEX2-ePTS1 construct before and after treatment with 10 µM MG132 for 4 h.
Areas presented in the figure are indicated.

10 % SDS PAGE. Stain-Free total protein staining was used as a loading control. The membrane was cut onto two parts. Upper part was used to detect RRBP1. Lower part was used for the detection of PEX3. Then the lower part was cut onto two parts and PEX16 was detected.

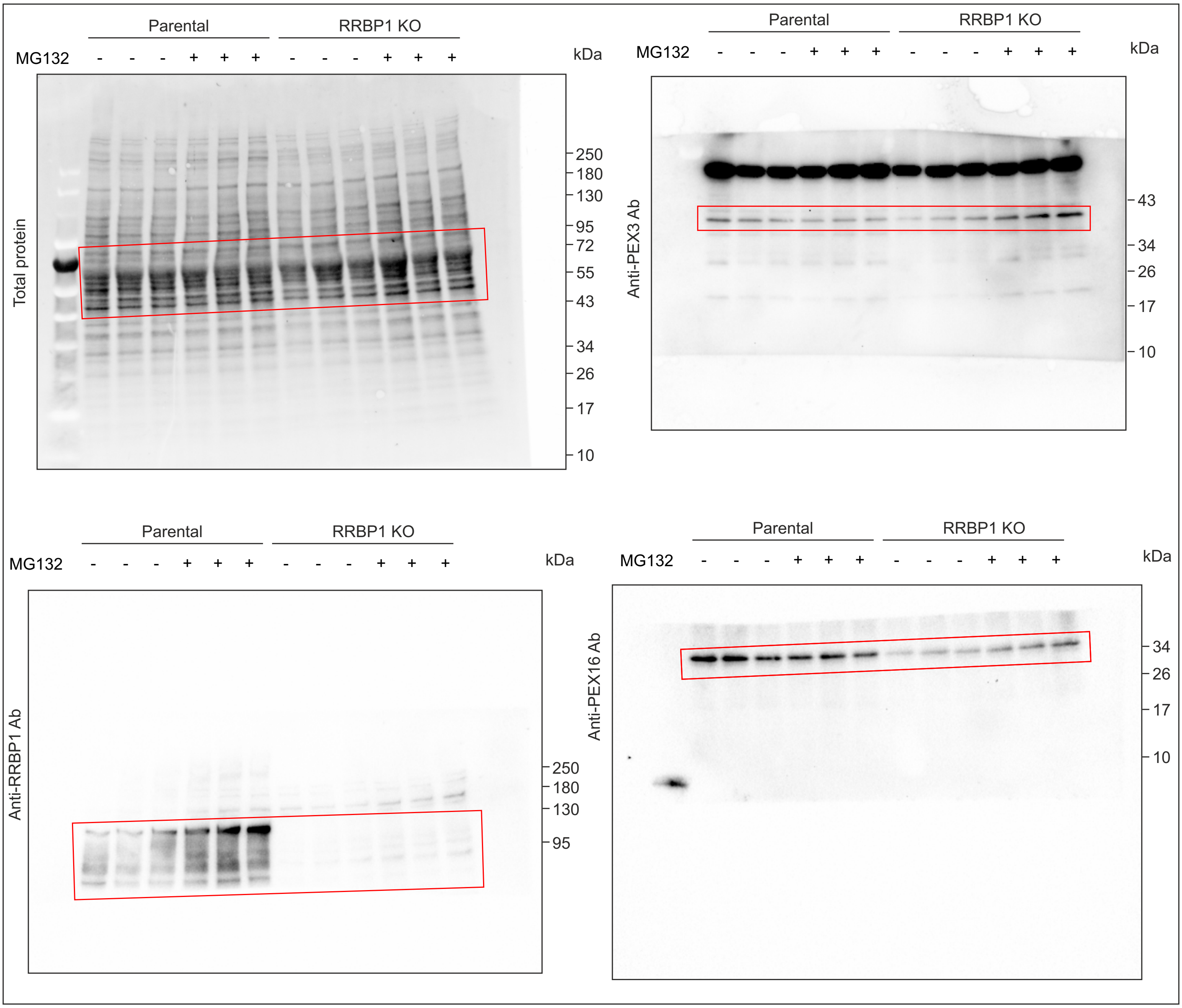

#### Unprocessed western blot images related to Supplementary Figure 1A

Western blotting analysis of HEK293T single cell clones stably expressing GFP-APEX2-ePTS1 or GFP-APEX2 constructs.
Areas presented in the figure are indicated.

Blot 1: 10% SDS PAGE. PEX19, GFP and GAPDH were detected on the same blot in the folllowing order: PEX19, GAPDH and GFP last.
Blot 2: 10% SDS PAGE. PEX3 and tubulin were detected on the same blot in the folllowing order: PEX3 first and then tubulin.

**Blot 1**

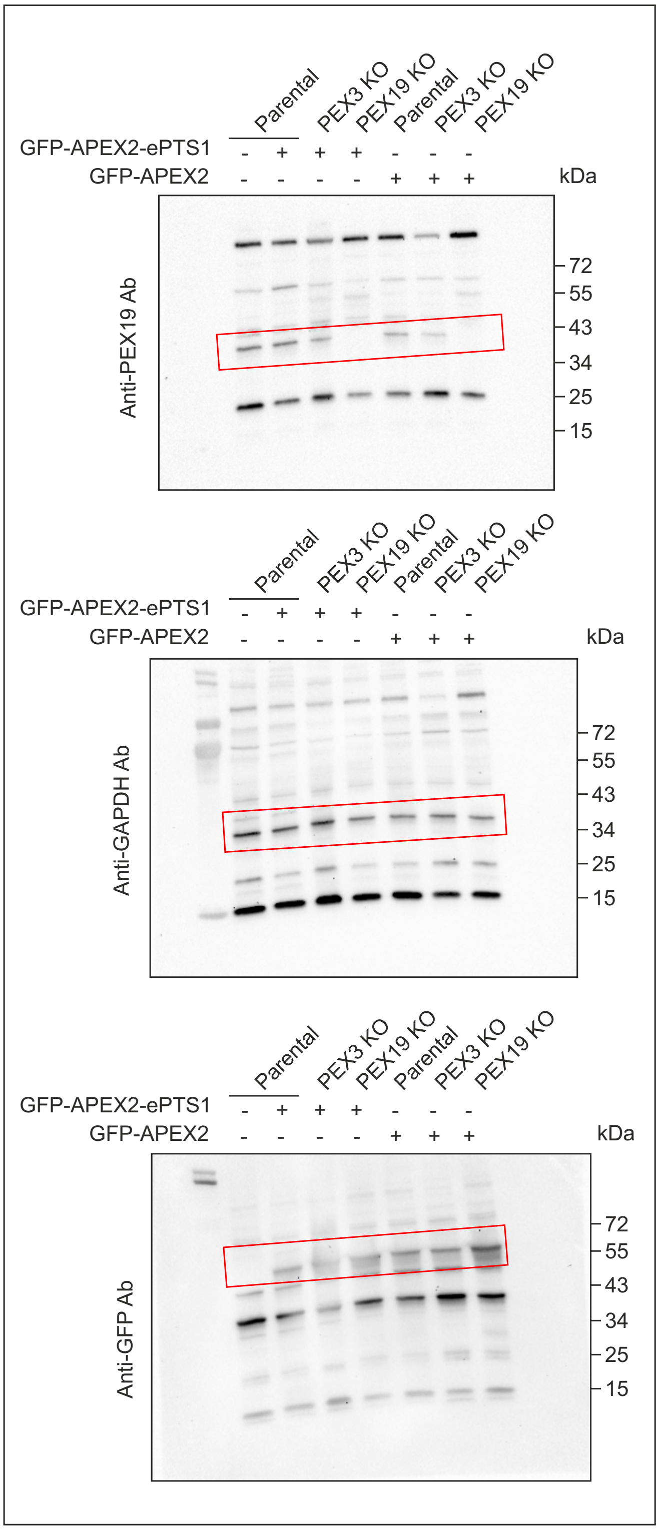


**Blot 2**

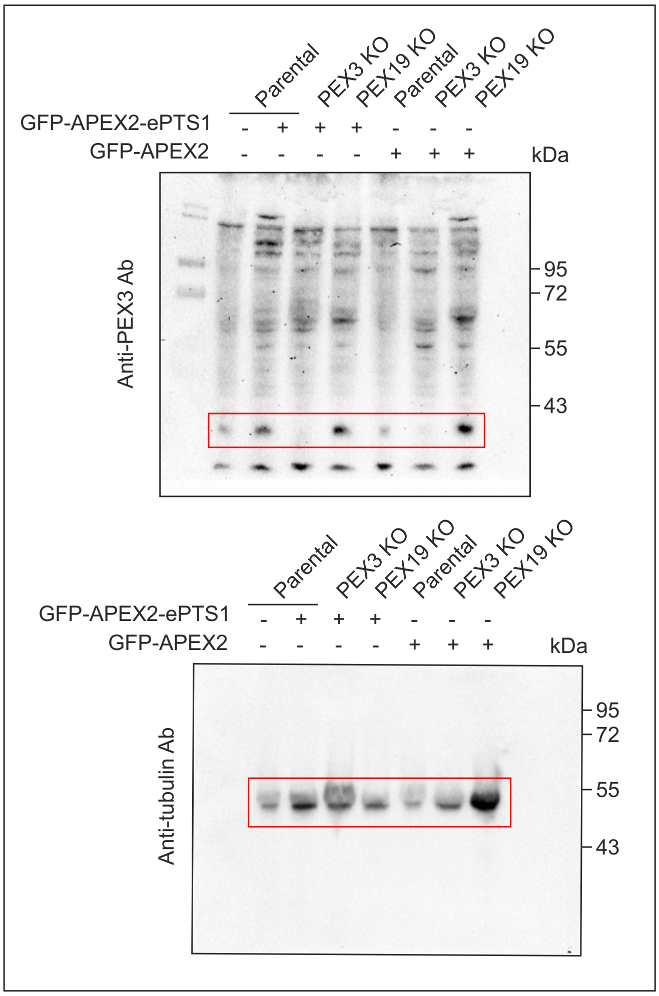

Blot 3: 10% SDS PAGE. PEX5 and GAPDH were detected on the same blot in the folllowing order: PEX5 first and then GAPDH.

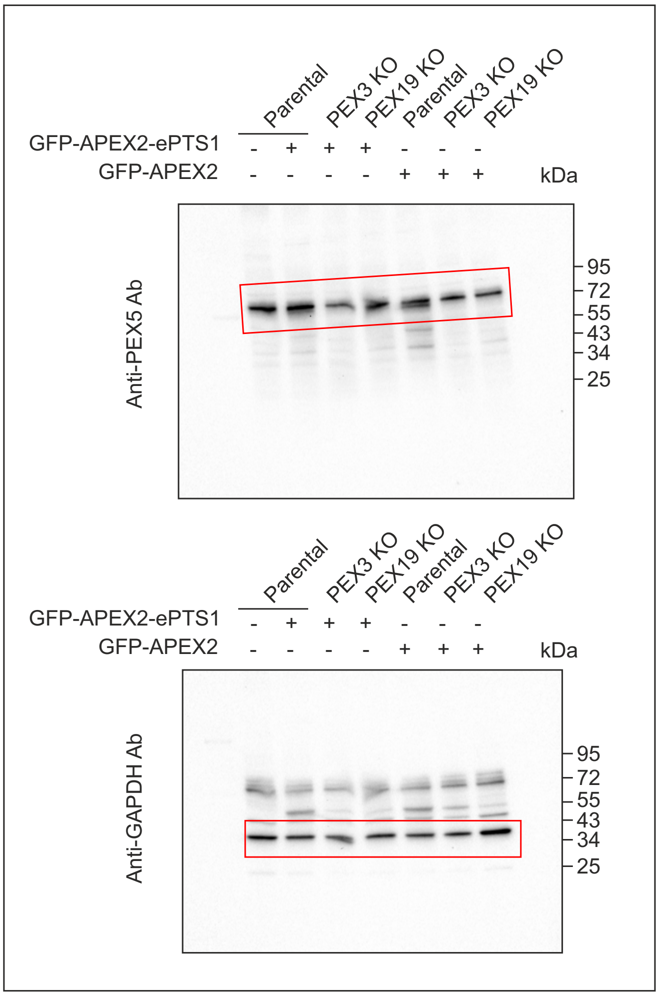

### Unprocessed western blot images related to Supplementary Figure 1E

Western blot analysis of GFP-APEX2-ePTS protein level in HEK293T cells stably expressing GFP-APEX2-ePTS1 before and after 13 days of transfection with Cas9-mCherry plasmid and gRNA targeting PEX19, PEX5 or RRBP1. Anti-GFP antibody was used to detect the GFP-APEX2-ePTS protein.
Areas presented in the figure are indicated.

Blot 1: 10% SDS PAGE. GFP and GAPDH were detected on the same blot.
Blot 2: 10% SDS PAGE. GFP and GAPDH were detected on the same blot.

**Blot 1**

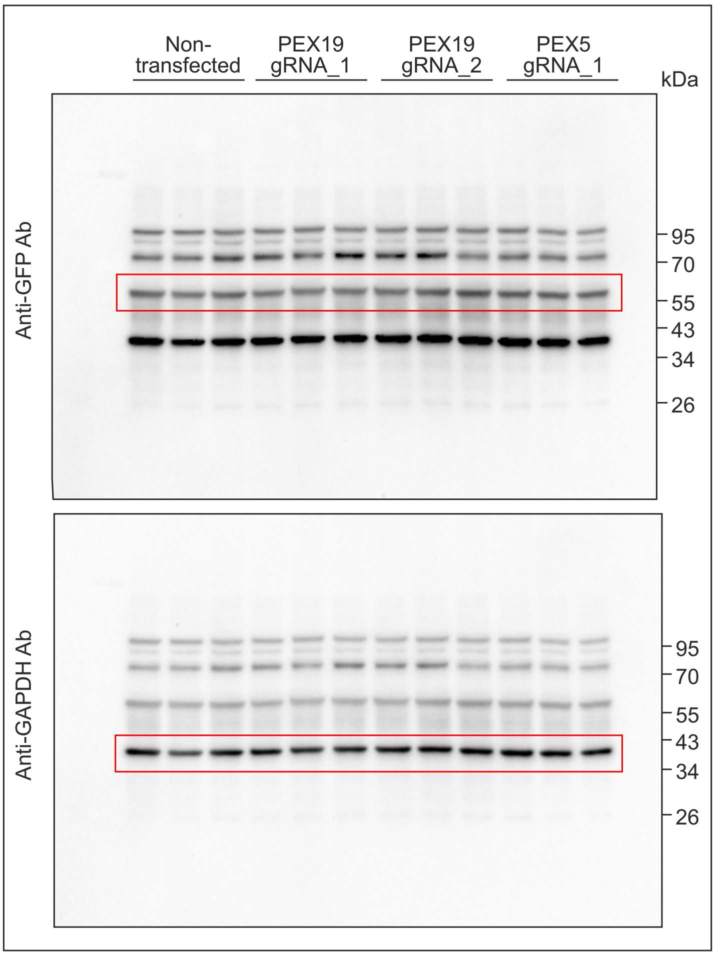


**Blot 2**

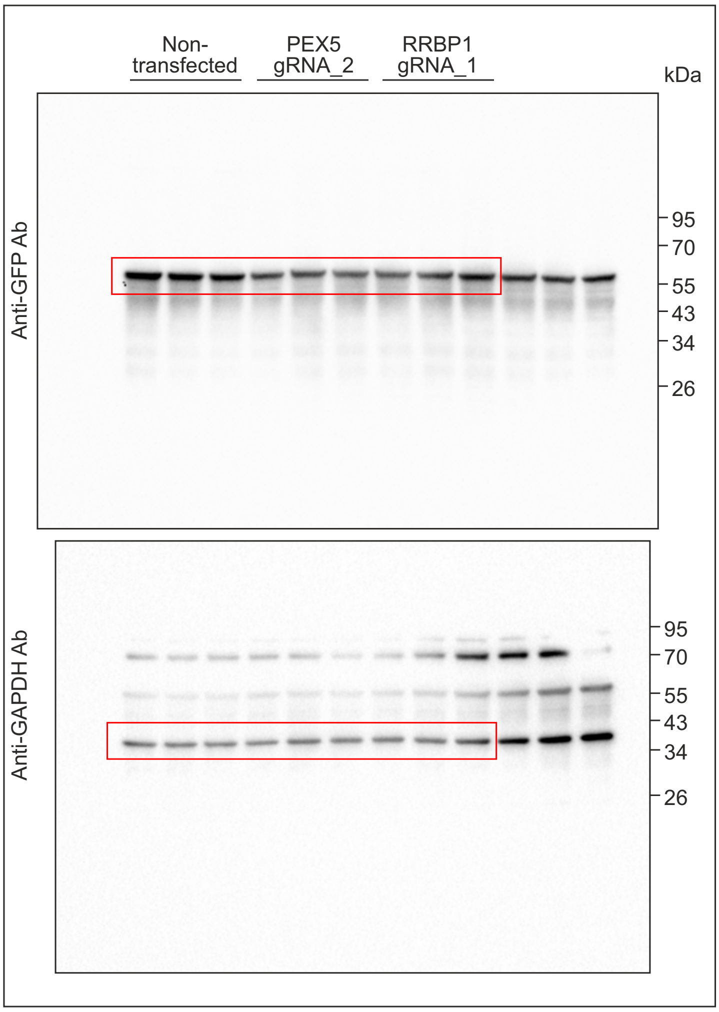

#### Unprocessed western blot images related to Supplementary Figure 2B

Western blot analysis of RRBP1 KO or parental HEK293T cells stably expressing GFP-APEX2-ePTS1 before or after puromycin treatment (10 µg/ml, 20 min).
Areas presented in the figure are indicated.

10% SDS PAGE. Puomycin and GAPDH were detected on the same blot.

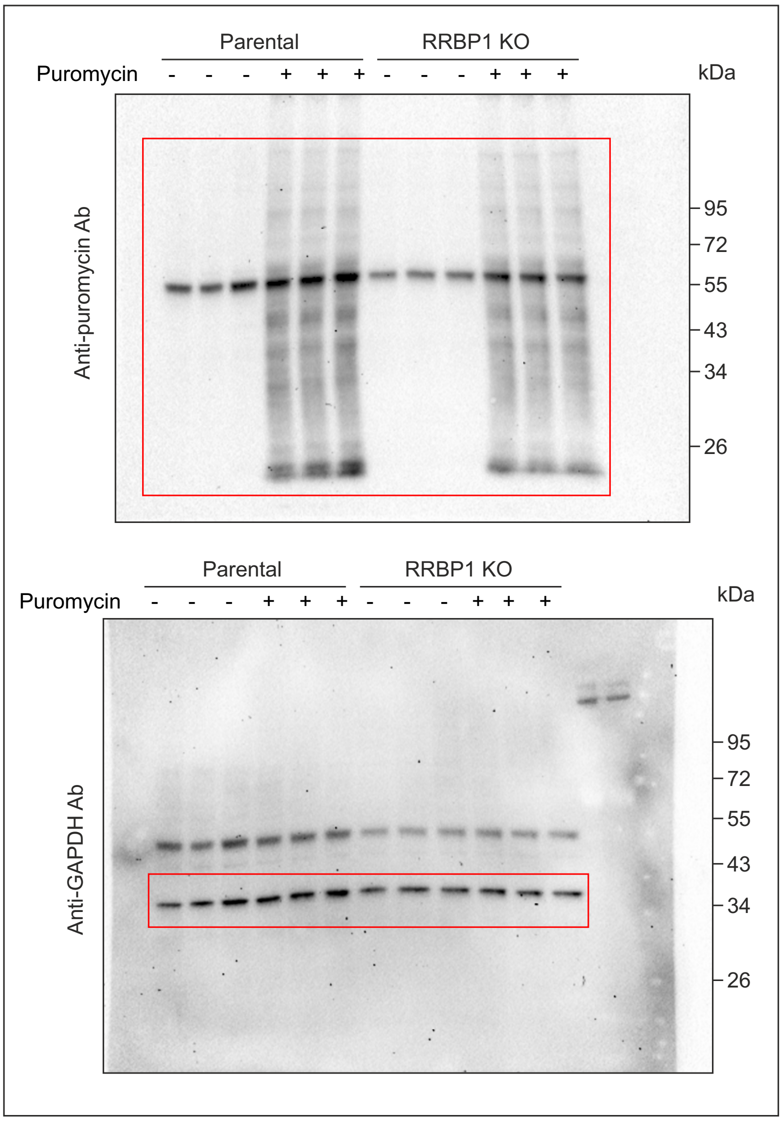

### Unprocessed western blot images related to Supplementary Figure 2D

Western blot analysis of RRBP1 KO HEK293T cells stably expressing GFP-APEX2-ePTS1.
Parental, PEX3 and PEX19 KO HEK293T cells not expressing peroxisomal targeted construct were used as controls. GAPDH was used as a loading control.
Areas presented in the figure are indicated.

Blot 1: 10% SDS PAGE. SEC61, IRE1, PXMP2 and GAPDH were detected on the same blot.

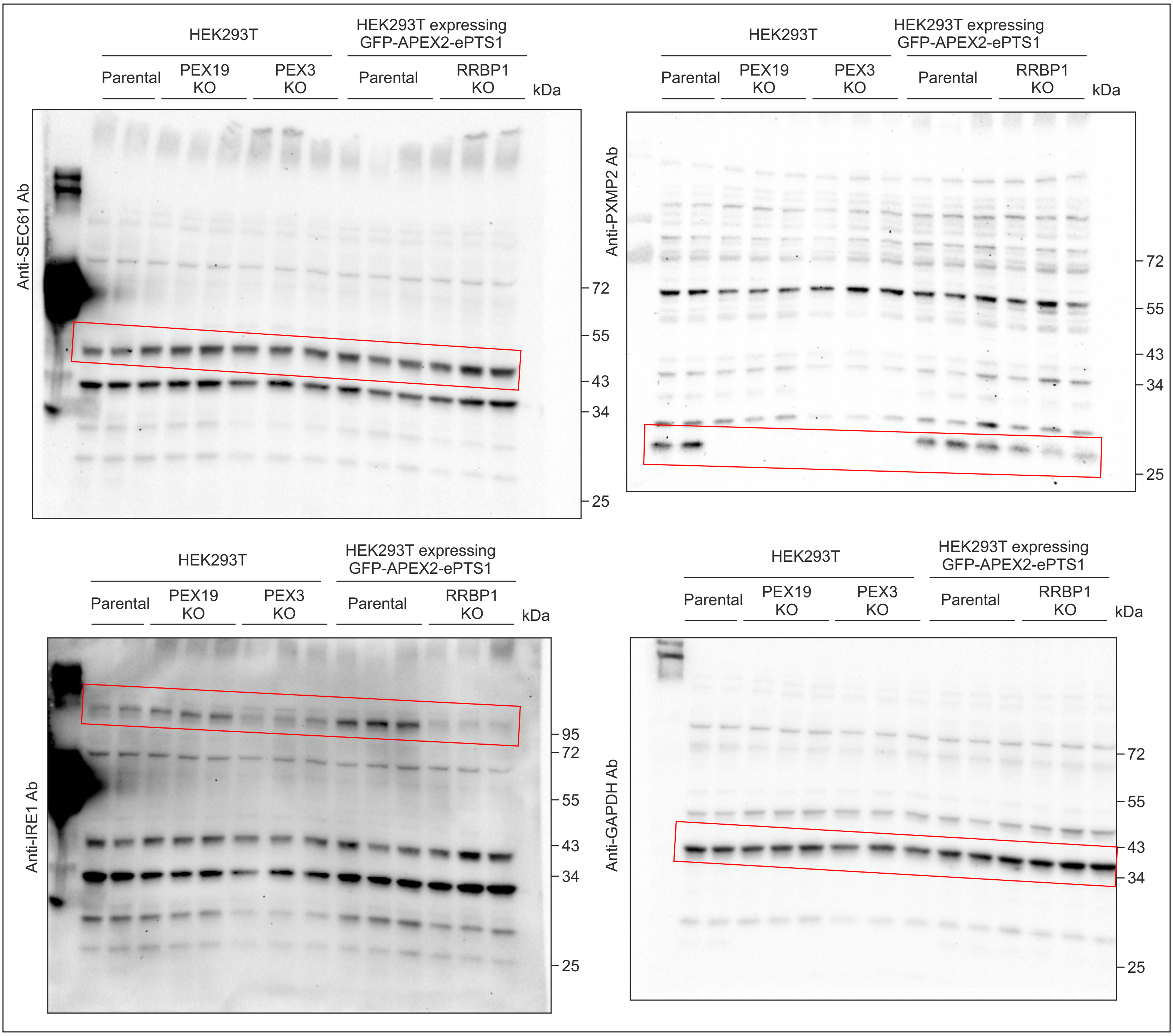

Blot 2: 10% SDS PAGE. BiP, RPL7 and GAPDH were detected on the same blot.

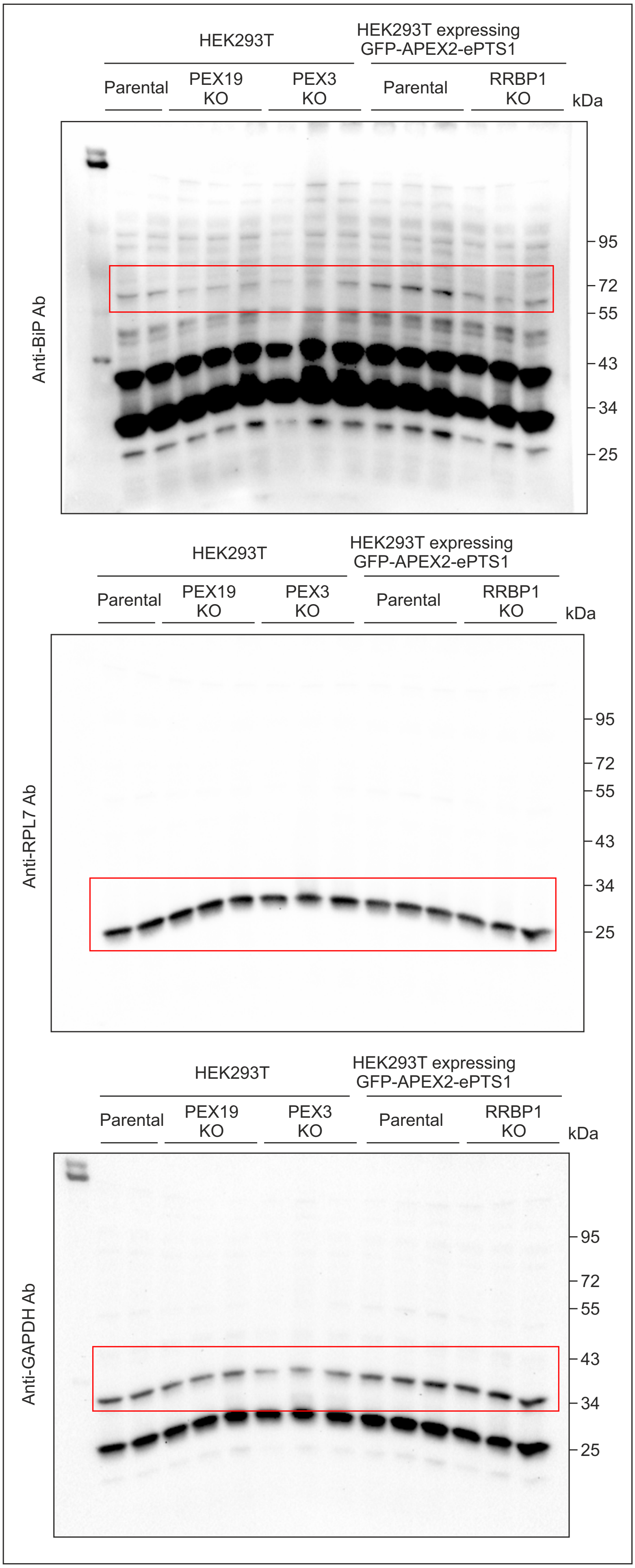
